# Targeting synthetic lethality between non-homologous end joining and radiation in very high-risk SHH medulloblastoma

**DOI:** 10.1101/2024.11.11.622104

**Authors:** Alexandria DeCarlo, Graham MacLeod, Carolina Fernandes da Silva, Lily Shen, Lucas Aragao, Mariska Sie, Deborah Termini, Jon Magee, Brian Gudenas, Sneha Sukumaran, Frederic Charron, Richard Marcellus, Rima Al-Awar, Ahmed Aman, Uri Tabori, Carolina Nör, Paul A. Northcott, Peter Dirks, Stephane Angers, Vijay Ramaswamy

## Abstract

Specific and biologically informed treatments for medulloblastoma, especially the highly lethal *TP53* mutant SHH subgroup, remain elusive, where radiotherapy is the primary treatment option. Applying genome-wide CRISPR-Cas9 screening in combination with lethal doses of radiotherapy, we identified the main driver of radiation resistance in SHH medulloblastoma is loss of p53. A negative selection CRISPR-Cas9 screen across multiple models of *Trp53*-deficient SHH medulloblastoma revealed a strong synthetic lethal interaction between components of the non-homologous end-joining pathway and radiation, particularly DNA protein kinase (DNA-PK) and its binding partners. Both genetic and pharmacological perturbation of DNA-PK enhanced radiosensitivity in *TP53*-deficient SHH medulloblastoma, leading to cell death. *In vivo* treatment of somatic and germline *TP53*-mutant SHH medulloblastoma models with peposertib, a small-molecule inhibitor of DNA-PK, significantly improved survival when combined with radiotherapy, strongly supporting further clinical investigation.

## Introduction

Medulloblastoma is the most common malignant brain tumor of childhood, and is comprised of four core molecular subgroups termed WNT, SHH, Group 3 and Group 4, each with distinct demographics, genetics and outcomes.^1–3^ Current therapy which dates back to the 1980’s is multimodal, involving aggressive surgical resections, craniospinal irradiation, and cytotoxic chemotherapy, resulting in five-year survival rates of approximately 50-60% for metastatic disease and 80% for non-metastatic disease.^4–6^ Radiotherapy remains the most effective treatment for medulloblastoma and has been the cornerstone of therapy since the early 1950s, with chemotherapy introduced in the 1980s providing marginal improvements in outcomes.^7,8^ Except for WNT medulloblastoma, which has been prioritized for clinical trials to reduce treatment intensity due to high survival rates, therapy is not tailored to the underlying molecular subgroup.^9^ A substantial proportion of patients are overtreated, however the subset of patients who relapse has remained consistent despite increased doses of radiotherapy and chemotherapy. The long-term side effects of this aggressive therapy on the developing brain are increasingly recognized, leaving most survivors with lifelong disabilities, including stroke, hearing loss, growth failure, hormonal deficiencies and cognitive impairment, highlighting the need for new treatment approaches.^10^

Clinical stratification currently lacks a biological basis and does not prioritize new therapies for patients with the worst prognosis. Advances in molecular characterization have significantly improved our understanding of the biological heterogeneity across medulloblastoma’s, clearly identifying some tumors as very high-risk with extremely poor outcomes.^9^ One such high-risk group is *TP53*-mutant SHH medulloblastoma (SHH-MB), which comprises a substantial portion of relapses and deaths in childhood. ^3,11^ *TP53* mutations in SHH-MB can occur both as somatic mutations and as germline mutations in the context of Li-Fraumeni syndrome. These tumors are characterized by genomic instability, frequent amplification of *MYCN* and/or *GLI2*, aneuploidy, and chromothripsis.^12,13^ Survival rates for both somatic and germline *TP53*-mutant SHH-MB are dismal, with overall survival of below 20% in most cohorts.^5,6,14,15^ SHH-MB tumors often relapse locally in the surgical cavity despite high doses of radiation, including a boost to the local tumor bed to 55.8 Gy, in contrast to Group 3 and Group 4 tumors, which almost exclusively relapse in the metastatic compartment.^16,17^ This suggests that radiation resistance is a significant cause of relapse and death in *TP53*-mutant SHH-MB. Furthermore, the few survivors of germline *TP53*-mutant SHH-MB almost always develop a second malignancy as a result of their therapy, leading to very poor long-term survival.^14,18^ *TP53* mutations can also emerge at relapse, yet clear mediators of radiation resistance in medulloblastoma, particularly SHH-MB, have yet to be identified and validated.^19,20^

Genome sequencing of SHH-MB has revealed a highly unstable genome many somatic copy number alterations, including focal amplifications and deletions, however there are a paucity of somatic nucleotide variants and actionable alterations.^13^ Most cases involve dominant negative mutations with loss of 17p, while a subset has *TP53* deletions. SHH pathway activation typically occurs downstream through *MYCN* and/or *GLI2* amplification, rendering SHH pathway inhibitors like vismodegib ineffective.^21^ Preclinical approaches to targeting *TP53*-mutant SHH-MB have been limited, and the only open clinical trial specifically for *TP53*-mutant SHH-MB, PNET5, focuses on omitting alkylator therapy and limiting the field of radiotherapy, rather than introducing targeted compounds.^22,23^

Radiotherapy is by far the most effective therapy for medulloblastoma.^1^ Despite the high relapse rates in very high-risk groups, including *TP53*-mutant SHH-MB, survival is significantly extended in radiated compared to non-irradiated patients.^14^ In the absence of clear actionable targets, one very attractive option is to pursue rational radiosensitizers, which have the potential to both increase the efficacy of radiotherapy, while also offering the potential to reduce radiation doses to patients with favourable outcomes, such as average risk *TP53*-wildtype SHH-MB.^5,6,9,11^ Although the children’s oncology group has conducted large studies of chemotherapy as a radiosensitizer, these studies have shown no benefit for the *TP53-*mutant SHH-MB subset of patients.^4,24^

In this study, we applied genome-wide CRISPR-Cas9 positive and negative screening to discover that the loss of p53 is the major determinant of radiation resistance in SHH-MB. Notably, we observed that dropout of key mediators of the non-homologous end joining (NHEJ) pathway, specifically DNA-protein kinase (DNA-PK) and its binding partners, are the only synthetic lethal interactions with radiotherapy. Inhibition of DNA-PK using peposertib, a potent and selective small-molecule inhibitor of DNA-PK, effectively radiosensitizes patient-derived orthotopic models of both somtic and germline *TP53*-mutant SHH-MB. Our results provide compelling evidence that DNA-PK inhibition is an effective therapeutic strategy to radiosensitize very high-risk *TP53*-mutant SHH-MB and warrants further investigation.

## Results

### p53 Deficiency Confers Radiation Resistance in SHH-MB

To understand the role of p53 status and radiation response in SHH-MB, we generated radiation dose-response curves from cell lines derived from four different *Ptch+/-;Trp53+/-het* (SHH*trp53*) tumors and two *Ptch+/-;Trp53+/+* (SHHwt 1) tumors (**Figure 1A**).^25^ We observe that SHH*trp53* models are significantly more resistant to radiotherapy than SHHwt models, whereby a lethal dose in SHHwt models was determined to be 4Gy compared to over 8Gy in our SHH*trp53* (**Figure 1A**).

**Figure 1:**
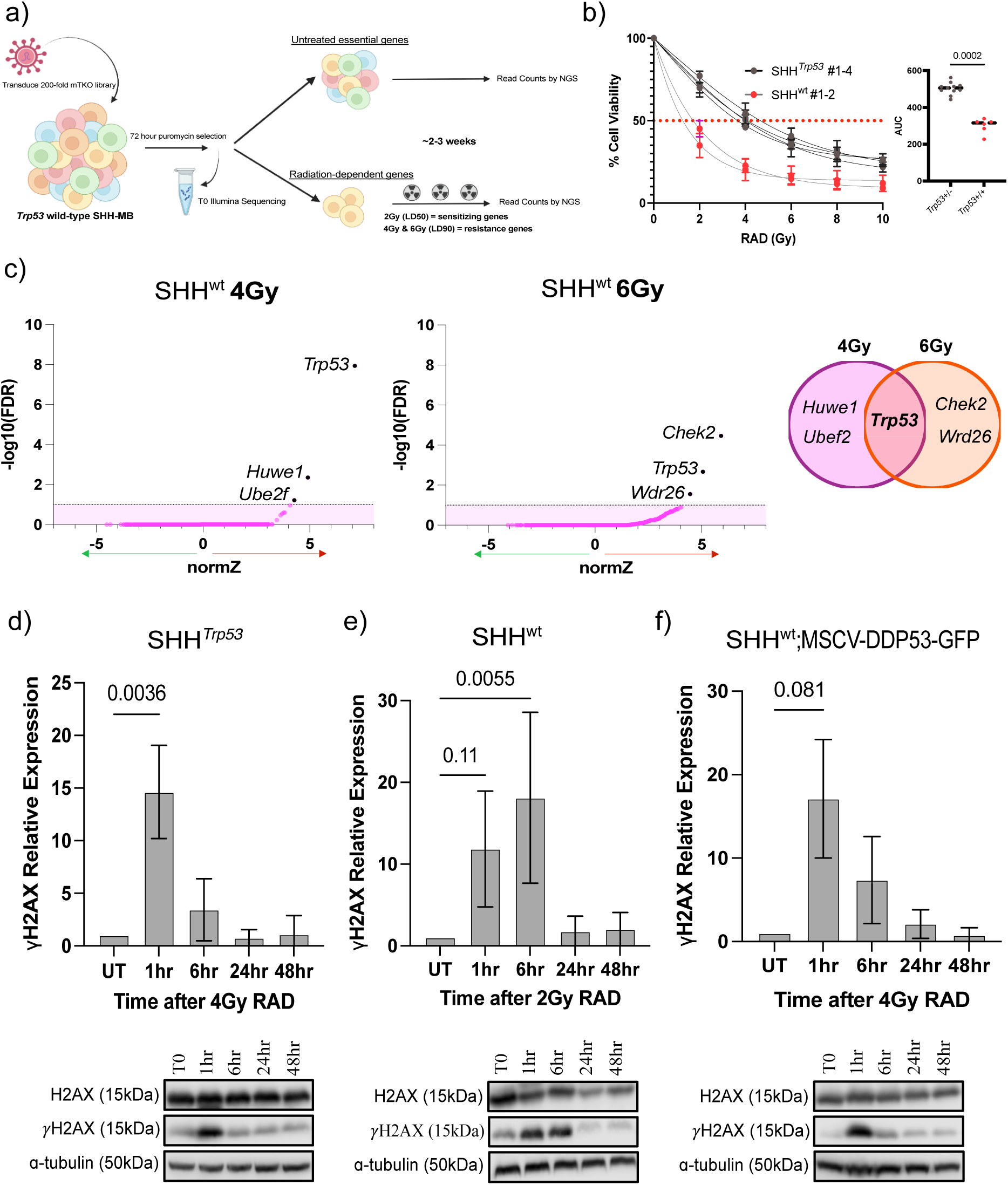
p53 deficiency confers radiation resistance in SHH-MB. **a)** Radiation dose-response curve (left) across a panel of SHH*trp53* (grey) and SHHwt (red). Cell viability was measured using alamarBlue and normalized to 0Gy and a positive control (puromycin). Analysis of the AUC from a non-linear regression of all models and replicates (right) demonstrates significantly increased (p=0.0002) radiation resistance in SHH*trp53*. Data was analyzed by Mann-Whitney test. **b)** Screen methodology. The mTKO library contains 94,528 gRNAs which target 19,069 protein-coding genes and are analogous to the human TKO version 3 (TKOv3) library, providing full coverage of the mouse genome. This screen is performed in three experimental arms with three technical replicates per arm, comparing no radiation with a sublethal and lethal dose of radiation to identify context-specific genes in SHH MB. Figure generated in biorender.com. **c)** Volcano plots of NormZ scores of gene essentiality versus -log10(FDR) in SHHwt treated with lethal doses of radiation. A positive NormZ score infers genes that cause resistance to radiotherapy. The shaded area represents the 10% FDR. **d-f)** Western blot analysis of yH2AX expression following radiation between SHH*trp53* **(e),** SHHwt **(f)**, and isogenic model **(g)** of SHH-MB over 48 hours. Data was normalized to protein loading control and T0 and analyzed by one-way ANOVA with Dunnett’s test for multiple comparisons to assess relative expression levels.

In order to elucidate potential mechanisms of radiation resistance, we performed two separate positive selection genome-wide CRISPR-Cas9 screens in our SHHwt model whereby we treated cells with lethal doses of 4Gy and 6Gy of radiation following transduction with the mouse Toronto knockout (mTKO) library (**Figure 1B, Figure S1A**).^26^ Integrated gRNAs within the remaining resistant population were sequenced and strikingly we found only three genes whose gRNAs showed significant enrichment within a 10% FDR threshold in both the 4Gy and 6Gy experiment. *Trp53* (p=5.94×10^-13^, FDR=1.6×10^-08^, normZ=7.11) was top-ranked and remained as the the only overlapping significant gene between the two screens (**Figure 1C, Table S1-2**). We then introduced a retrovirus with the *Trp53*^DD^ allele^27^ into the SHHwt 1 model to create an isogenic model of SHH*trp53* tumors, and we observe that p53-deficiency significantly increases radiation resistance compared to our SHHwt model (**Figure S1B**). *Ptch+/-;Trp53-/-* (null) and DAOY p53-mutant human medulloblastoma showed similar radiation dose response curves as the SHH*trp53* models (**Figure S1C-S1D**). Low pass whole genome sequencing of the four SHH*trp53* models showed numerous copy number changes reminiscent of human *TP53*-mutant SHH-MB, however introduction of the *Trp53*^DD^ allele did not, suggesting that *Trp53*-deficiency alone is sufficient to cause radiation resistance in SHH-MB (**Figure S1E -S1F**).

In order to evaluate DNA-repair capacity in our models, we proceeded to measure levels of γH2AX, a marker of DNA double-strand breaks (DSBs), across our models over 48 hours following exposure to radiation (**Figures 1D-F**). All models showed increased expression of γH2AX 1 hour following radiation, however, SHHwt showed increased expression at the 6-hour timepoint whereas SHH*trp53*, including our dominant negative isogenic model, returned to baseline expression by 6 hours. This suggests that in the absence of p53 function, DNA repair capacity may be attenuated compared to wild-type p53, consistent with the known functions of p53. Taken together, p53 deficiency alone confers radiation resistance in SHH-MB and influences the capacity for DSB repair.

### Genome-Wide CRISPR-Cas9 Screening Identifies Modulators of Radiation Sensitivity in *Trp53*-Deficient SHH-MB

Having identified that p53-defiency confers resistance to radiotherapy, and results in an attenuated DNA-damage response, we asked if this could be leveraged to identify synthetic lethal interactions which can serve as targets for radiosensitization. To identify genes which may sensitize SHH*trp53* to radiation, we performed negative selection genome-wide CRISPR-Cas9 screens across three SHH*trp53* models and one SHHwt model (**Figure 2A**).^28^ SHH*trp53* models were treated with a sublethal dose (4Gy) of radiation following transduction of the mTKO library, and allowed to propagate 14 doublings. Samples were collected after several cell doublings following radiation exposure and sequenced to analyse for depleted gRNAs whereby their perturbations would sensitize the cells to radiation.^29^ NormZ scores were used to identify synthetic lethality across our models whereby any hits with a negative NormZ score within the FDR threshold of 10% are ideal targets for radiation sensitization. Multiple genes confer sensitization across all models and demonstrate several pathways, with the most prominent genes being involved in DNA break repair (**Figure 2B**). A lethal dose of 8Gy or 12 Gy (4Gy x 3 doses) was also applied to all three SHH*trp53* models to look for genes conferring resistance (positive NormZ score) and no common genes or pathways were identified **Figure S2A-C**.

**Figure 2:**
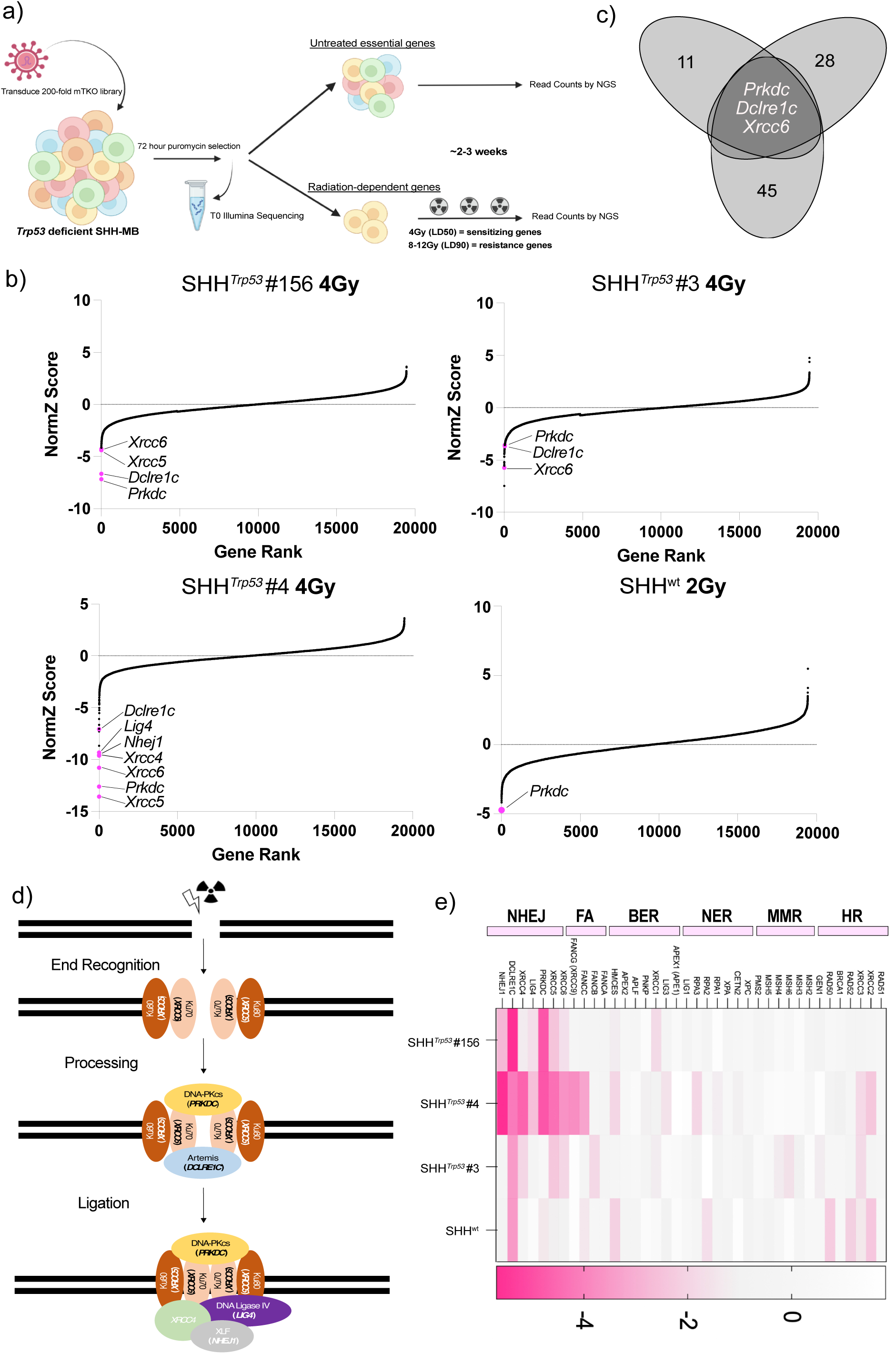
*Trp53* deficient SHH-MB rely on NHEJ following radiation. **a)** Screen methodology as described in Figure 1a. Figure generated in biorender.com. **b)** NormZ scores at midpoint across all screened SHH*trp53* models and one SHHwt one with genes highlighted in pink being involved in NHEJ and below the FDR threshold of 0.1 (10%). **c)** Venn Diagram of normZ score of -1 or less with 10% FDR threshold at midpoint, representing essential genes for response to radiation across all three SHH*trp53*, highlighting key three genes involved in NHEJ. **d)** Schematic diagram of NHEJ DNA DSB repair. Genes and the corresponding protein highlighted in black represent enriched targets from the CRISPR screen. **e)** Heatmap of quantile normalization of NormZ scores at endpoint divided into different DNA repair pathways across both *Trp53* deficient and wildtype SHH-MB models.

Although each individual screen resulted in several genes contributing to radiation sensitization within the FDR threshold of 10% (**Tables S3-10**), when comparing the common genes across all three models, only three remain (**Figure 2C**). *PRKDC*, encoding for DNA-PK, *DCLRE1C*, encoding for artemis, and *XRCC6*, encoding for Ku70, one-half of the Ku70/80 heterodimer. All three of these genes encode for essential proteins involved in non-homologous end-joining (NHEJ), a major pathway for DSB repair.^30^ Interestingly, within each screen, most genes involved in NHEJ were depleted across all SHH*trp53* models, as indicated by their negative normZ score (**Figure 2B, 2D**). In the SHHwt model treated with a sublethal dose of radiation, we only observed enrichment of *PRKDC,* which was then lost by the end of the screen, without enrichment of other components of the NHEJ pathway (**Figure 2D, Table S9-S10**). Surprisingly, no common hits were identified in homologous recombination (HR), another major repair pathway for DSBs, across our models. We further analyzed our data through quantile normalization of the NormZ scores across all models screened at endpoint, including the SHHwt model (**Figure 2E**). When contrasting synthetic lethal interactions with radiotherapy across different DNA repair pathways, we find a clear enrichment for NHEJ-related genes across SHH*trp53*. Taken together, this data suggests that although SHHwt may have some sensitivity to disruption of *PRKDC* (**Figure S3B**), the absence of p53 function may lead to a unique dependency on NHEJ in SHH-MB following radiotherapy, likely secondary to their decreased capacity for DNA-repair following DSBs, and suggests inhibiting NHEJ is a rational therapeutic vulnerability to functionally validate.

### Genetic and Therapeutic Targeting of DNA-PK and *PRKDC* Sensitizes SHH-MB to Radiation

*PRKDC* encodes for DNA protein kinase (DNA-PK), an essential kinase for DNA repair which coordinates end-processing and end-ligation during NHEJ through interactions with additional proteins involved in DSB repair.^31^ DNA-PK is thought to be the master regulator of NHEJ and is activated through either auto-phosphorylation or ATM-mediated transphosphorylation whereby autophosphorylation is critical for NHEJ function. Several small molecule inhibitors against DNA-PK have been developed that are both specific and potent however both artemis and the ku70/80 heterodimer are non-targetable, and as such DNA-PK was prioritized as a potential therapeutic target. To confirm that genetic knockout of DNA-PK is synthetically lethal with radiation, we generated a CRISPR-Cas9 single-gene knockout of *PRKDC* (**Figure 3A**) and confirm that this successfully sensitizes SHH*trp53* models to radiation while having no significant effect on cell fitness in non-radiated cells (**Figure 3B**).

**Figure 3:**
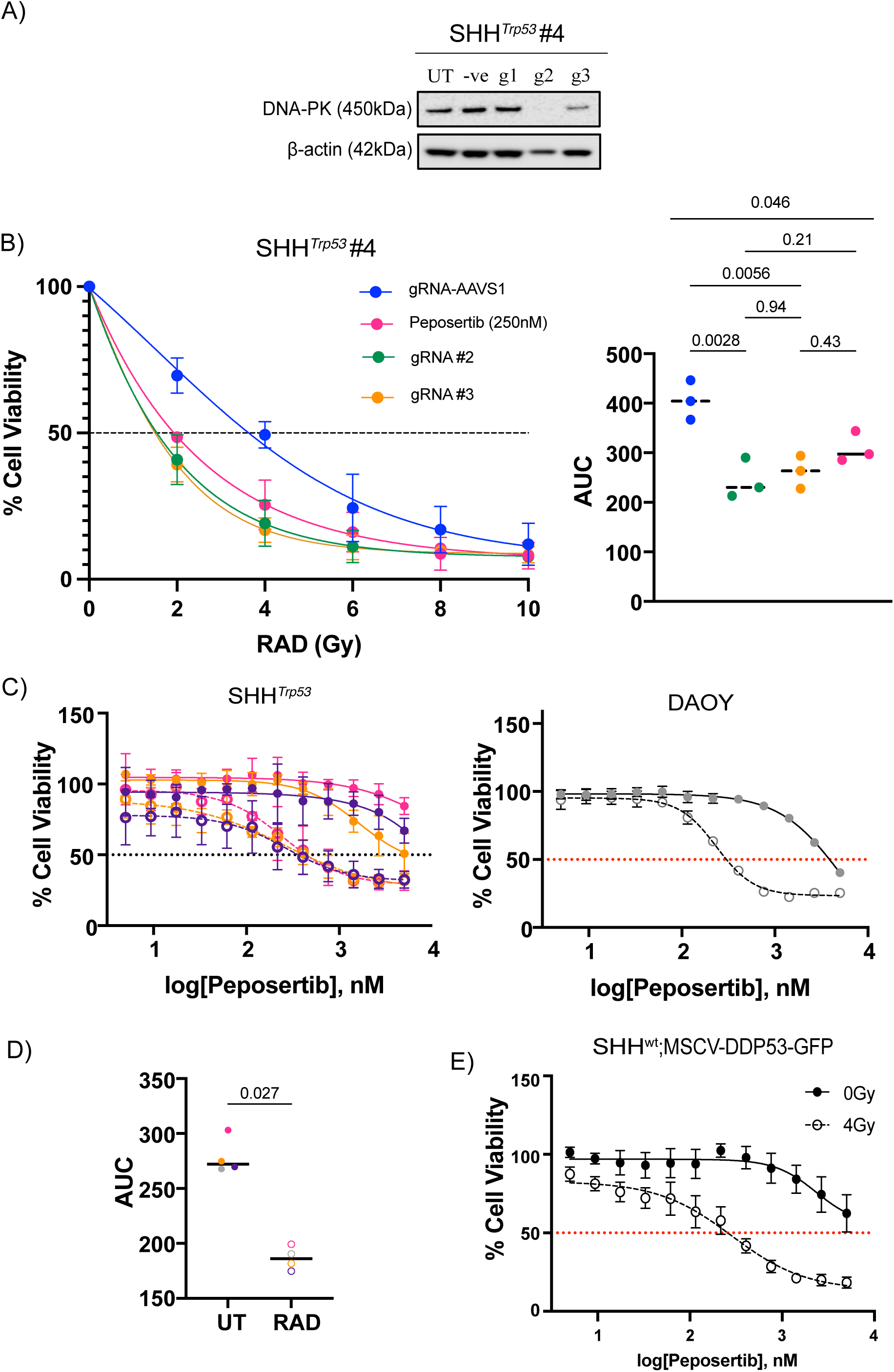
Genetic and therapeutic targeting of PRKDC and DNA-PK sensitizes *Trp53* deficient SHH-MB to radiation. **a)** Western blot demonstrating successful Crispr-Cas9 mediated single gRNA knockout of *PRKDC*. **b)** Radiation dose-response curve (left) testing radiation sensitivity between two PRKDC genetic knockouts (pink and orange), 250nM of peposertib (green), and a non-targeting control (blue). Analysis of the AUC from a non-linear regression of all replicates demonstrating a significant increase in sensitivity between the non-targeting control and gRNA#2 (p=0.0028), gRNA#3 (p=0.0056), and peposertib (p=0.046) with no significant difference in sensitivity between peposertib and gRNA#2 and gRNA#3 (p=0.21, 0.43, respectively). **c)** 12-point dose-response curves with peposertib with and without a sublethal dose (4 and 6Gy, respectively) of radiation for Trp53-decificnt SHH-MB (left) and Daoy (right). Solid colours represent no radiation and boardered colours represent sublethal radiation. **d)** Analysis of the AUC from a non-linear regression across all models demonstrating a significant (p=0.027) increase in sensitivity when treated with peposertib. **e)** 12-point dose-response curve with peposertib with and without a sublethal dose of radiation for SHH^wt^;MSCV-DDP53-GFP.

We next screened a panel of small-molecule inhibitors targeting DNA-PK. Here, we performed 12-point dose-response curves with and without radiation across several compounds and assessed for radiosensitization (**Figure S3A**). Both peposertib (also known as nedisertib) and AZD7648 displayed significant radiosensitization across three SHH*trp53* models, with minimal effect of the drug alone (**Figure 3C-D and S3A**). Furthermore, we observe peposertib is a potent radiosensitizer of the human SHH-MB model DAOY, and has a more potent ability to radiosensitize *Trp53*^DD^ transduced *ptch+/-* cells than *Trp53* intact (**Figures 3C-E and S3B**). We next performed Caco-2 permeability assays, an accurate indicator of which compounds are permeable based on their ability to move through tight junctions with minimal efflux^32^, with our compounds of interest and observe that peposertib is a stable blood-brain barrier permeable compound with minimal efflux (**Figure S3C-S3D**). Overall this suggests that the DNA-PK inhibitor peposertib is a potent radiosensitizer, with similar effects to knocking out *PRKDC*, confirming small molecule DNA-PK inhibition as a promising therapeutic target.

### Therapeutic Inhibition of DNA-PK Sustains Sensitization Over Several Cell Doublings

Within the context of radiation, effects are often not immediate and instead becomes obvious following several cell doublings through which cells acquire the accumulation of DNA damage. This ultimately results in cell death several days following the initial insult. Therefore, in order to fully understand the potential of DNA-PK inhibition as a radio-sensitizing strategy, we need to investigate the effects longer term. We treated SHH*trp53* (**Figures 4A and S4A**) and human DAOY cells (**Figure 4B**) with and without radiation and peposertib and monitored their growth over several cell doublings. Using IncuCyte Live Cell Imaging (**Figures 4A, S4A-S4B**), we monitored cell growth over 7 days following treatment of peposertib, radiation, or treatment combination.

**Figure 4:**
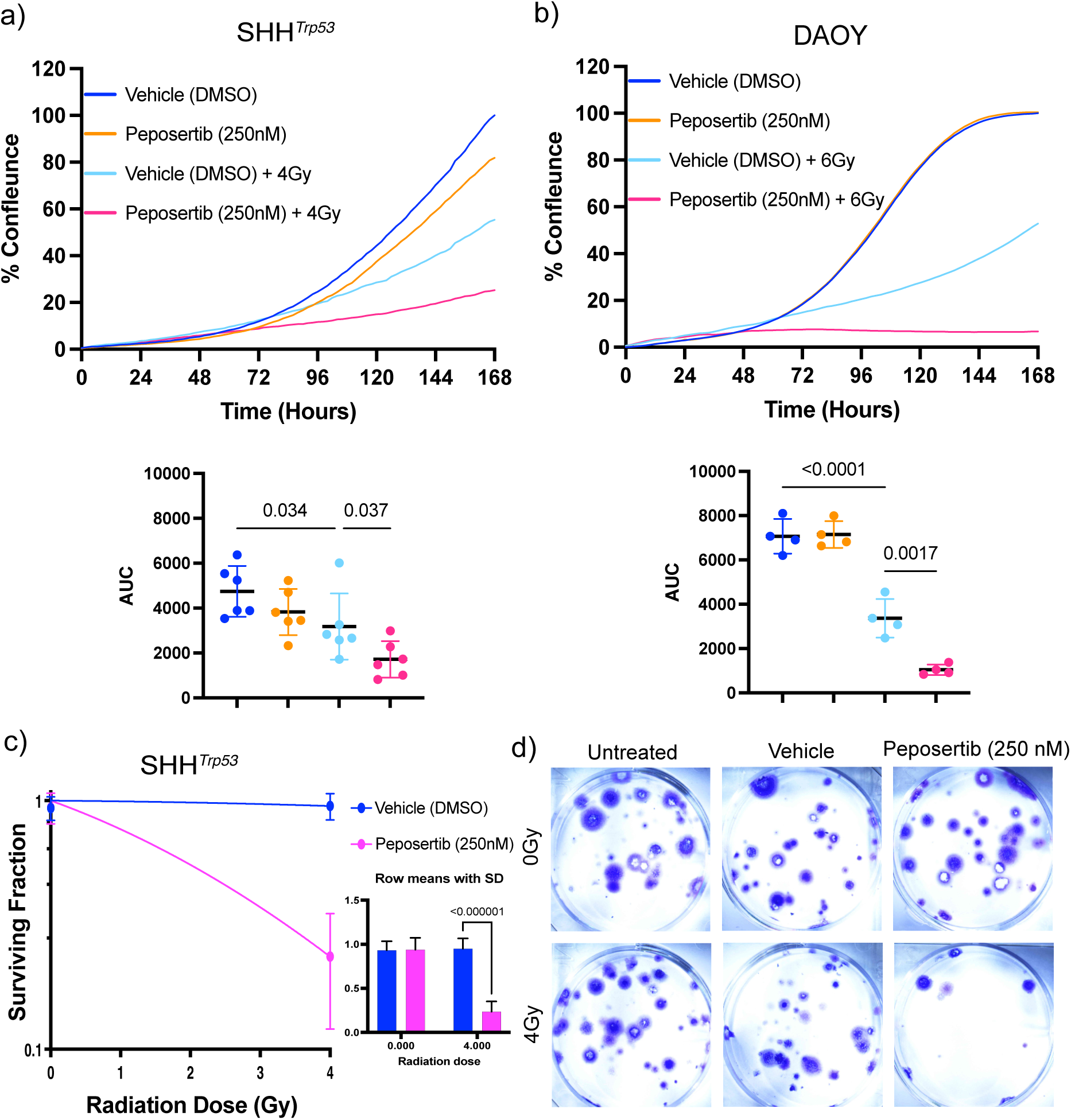
Therapeutic inhibition of DNA-PK sustains radiation sensitization over several cell doublings. **a)** Cell growth assays over 7 days with and without radiation in combination with 250nM of peposertib. Cell images were taken by Incucyte from four different fields/well every 2 hours over 7 consecutive days to infer cell growth. Cell confluence plotted as percent of the confluence determined in the DMSO controls (set as 100%). Top graph shows combined mean across multiple SHH*trp53* models. Bottom graph represents AUC of each independent assay, analyzed by one-way ANOVA with Tukey’s multiple comparisons test demonstrating a significant decrease in cell growth between radiation alone and radiation in combination with peposertib (p=0.037). **b)** Cell growth assay in human SHH-MB Daoy cells over 7 days as described above. Top graph shows combined mean across four replicates. Bottom graph represents AUC of each independent assay analyzed by one-way ANOVA with Tukey’s multiple comparisons test demonstrating a significant decrease in cell growth between radiation alone and radiation in combination with peposertib (p=0.0017). **c)** 14-day clonogenic assay combined analysis of multiple SHH*trp53* models with and without radiation in combination with 250nM of peposertib. Surviving fraction is calculated based on the number of colonies formed normalized to seeding density and plating efficiency. Data is analyzed with multiple unpaired t-tests of means of all replicates demonstrating significant decrease in clonogenic potential (p<0.0001) when cells are treated with a combination of radiation and peposertib. **d)** Representative images of clonogenic assays for one SHH*trp53* model.

Although radiation did slow cell growth, cells treated with radiation alone still had positive growth and were able to reach approximately 50% confluency over the 7 days, indicating their inherent resistance through their ability to continue to replicate. In contrast, when peposertib is combined with radiation cell confluence remains the same over the 7 days, appearing to plateau growth at 96 hours (**Figures 4A-4B)**. Peposertib alone had no effect on cell growth.

We then assessed clonogenic survival of SHH*trp53* cells in response to both peposertib and radiotherapy, and observe a significant reduction in clonogenic capacity in the combination therapy, whereas peposertib alone had no effect on colony formation (**Figure 4C-4D**). Taken together this suggests that the combination of radiotherapy and DNA-PK inhibition sensitizes cells through a combination of cell death and impaired long-term clonogenic survival, with no effect on non-radiated cells.

### Therapeutic Inhibition of DNA-PK Induces Necrosis in *Trp53*-Deficient SHH-MB

The DDR consists of both cell death and cell survival mechanisms to either stop cell cycle progression to facilitate DNA repair or to induce apoptosis or senescence if the damage is irrepearable. To further understand how the inhibition of DNA-PK influences the DDR to sensitize SHH-MB to radiation and to confirm our results from Figure 4, we looked at three essential effector pathways within the DDR. Utilizing flow cytometry for propidium iodide, annexin V and β-galactosidase, we investigated the percent of the cell population entering cell death, cell-cycle arrest, and senescence following radiation with or without peposertib over 96 hours.

The combination of peposertib and radiation resulted in a significant increase in cell death through necrosis compared to radiation alone by 96 hours following treatment (**Figure 5A-C**). Consistent with peposertib being a selective and potent radiosensitizer, peposertib alone had no effect on cell viability over 96 hours. Similarly, combinational treatment resulted in a shift from S-Phase to a significant increase G2 cell cycle arrest by 96 hours following treatment (**Figure S5B-C**) compared to vehicle but not to radiation alone. Additionally, we interpreted levels of senescence by measuring levels β-galactosidase. We found a slight increase in senescence in cells treated with combinational therapy at 96 hours (**Figure 5D**). Although there are increases in G2 cell cycle arrest and senescence, there is only a significant increase in necrosis between radiation alone and radiation plus peposertib. These results indicate that upon inhibition of DNA-PK, radiation induces DSBs that can no longer be repaired resulting in increased cell death thus sensitizing cells to radiation while having no effect on cells that are not treated with DNA-damaging agents.

**Figure 5:**
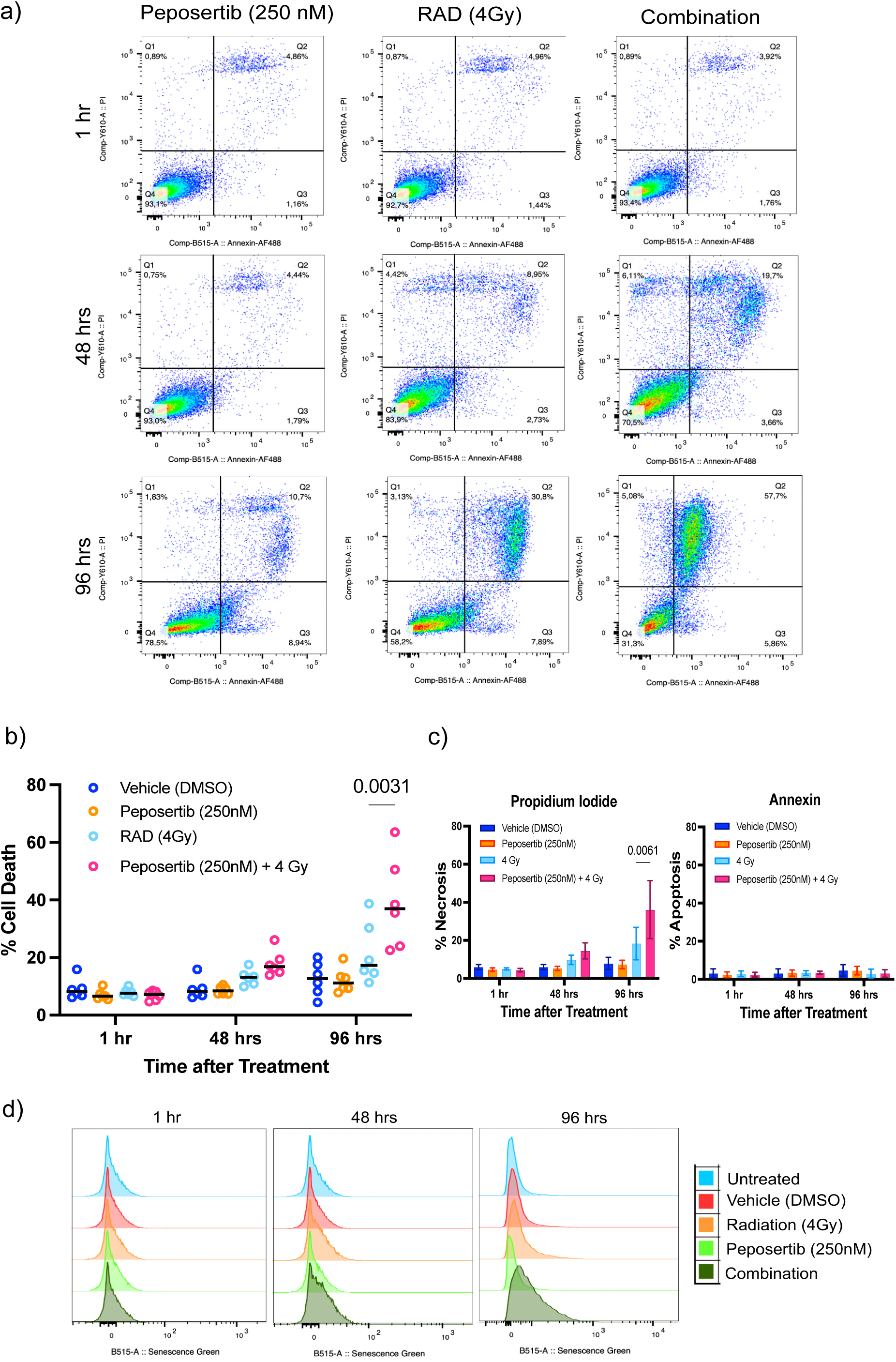
Therapeutic inhibition of DNA-PK induces necrosis in *Trp53* deficient SHH-MB. **a)** Representative images of Annexin V/PI flow cytometry cell viability assay in one SHH*trp53* model at multiple timepoints. **b)** Combined analysis of two SHH*trp53*models, each with three independent technical and biological replicates of Annexin V/PI assay at multiple timepoints representing total % cell death demonstrating a significant increase in cell death after 96 hours with combination therapy compared to radiation alone (p=0.0031). **c)** Combined analysis of two SHH*trp53* models, each with three independent technical and biological replicates of Annexin V/PI assay at multiple timepoints representing % cell death divided between apoptosis (right) and necrosis (left). Data analyzed by two-way ANOVA with the Geisser-Greenhouse correction with matched values and Tukey’s multiple comparisons test demonstrating a significant increase in necrosis with combination therapy compared to radiation alone (p=0.0061). **d)** Analysis of senescence with ß-galactosidase via flow cytometry in one SHH*trp53* model at multiple timepoints.

### Inhibition of DNA-PK with Peposertib results in potent radiosensitization *in vivo*

On the basis of our promising *in vitro* findings that both genetic and pharmacological inhibition of DNA-PK radiosensitizes p53 deficient SHH-MB, we further evaluated the combination of peposertib *in vivo*. Two p53 mutant SHH-MB patient derived orthotopic models were used, one somatic *TP53* mutant harboring a *MYCN*, *MDM4* and *CDK4* amplification (SJMBSHH -14-4106) and one germline *TP53* mutant harboring a *MYCN* amplification (SJMBSHH -13-5634). Cells were stereotactically implanted into the cerebella of 6-week old NRG (NOD rag gamma) mice, and 10 days after implantation animals were randomized to four arms: vehicle x 5 days, peposertib 50mg/kg twice daily x 5 days, 2Gy radiation daily x 5 days and peposertib 50mg/kg twice daily plus 2Gy radiation daily (**Figure 6A**). CT-guided radiotherapy was delivered to the brain and cerebellum. In the SJMBSHH-14-4106 model peposertib alone did not increase survival compared to vehicle, however the combination of peposertib and radiotherapy significantly increased survival compared to radiotherapy alone (**Figure 6B**, p<0.0001 Log-Rank, p=0.017 for only radiation alone vs combination therapy). Two animals in the combination arm (M4 and M5) had no tumor within the radiation field upon necropsy, but had metastatic disease along the spinal cord but not in the brain or cerebellum (**Figure 6C-D, and S6D-E**). One animal in the combination arm had in field failure (M1) and the remaining two animals remain alive and tumor free at 90 days post treatment. We treated an additional cohort of the SJMBSHH-14-4106 model with craniospinal irradiation in combination with or witout peposertib (**Figure S7A**) following tumor engraftement visualized by MRI. Again, we saw a significant increase in survival in our combination arm (**Figure S7B**, p=0.0016 Log-Rank). Here, we saw complete tumor reduction that was sustained compared the cohort of radiation alone, which all succumbed to local recurrence (**Figure S7C**). In the SJMBSHH-13-1-5634 model, there was also a significant survival benefit observed for the combination of peposertib and radiotherapy versus radiotherapy alone, with no survival benefit observed for peposertib alone (**Figure 7A**, p=0.0049 Log-Rank). Two animals in the combination therapy arm died of hydrocephalus 3 weeks after treatment with radiation and peposertib, and small amounts of tumor was identified, and the remaining three animals died of local failure significantly later than the animals in the radiation alone arm (**Figure 7B, 7C, S7E-F**). In the radiotherapy alone arm, all animals had visible tumor by both MRI and at necropsy (**Figure 7B, 7C**). Taken together, our results reveal that the combination of DNA-PK inhibition using peposertib and radiation significantly improves survival in PDX models of very high-risk *TP53*-mutant SHH medulloblastoma.

**Figure 6:**
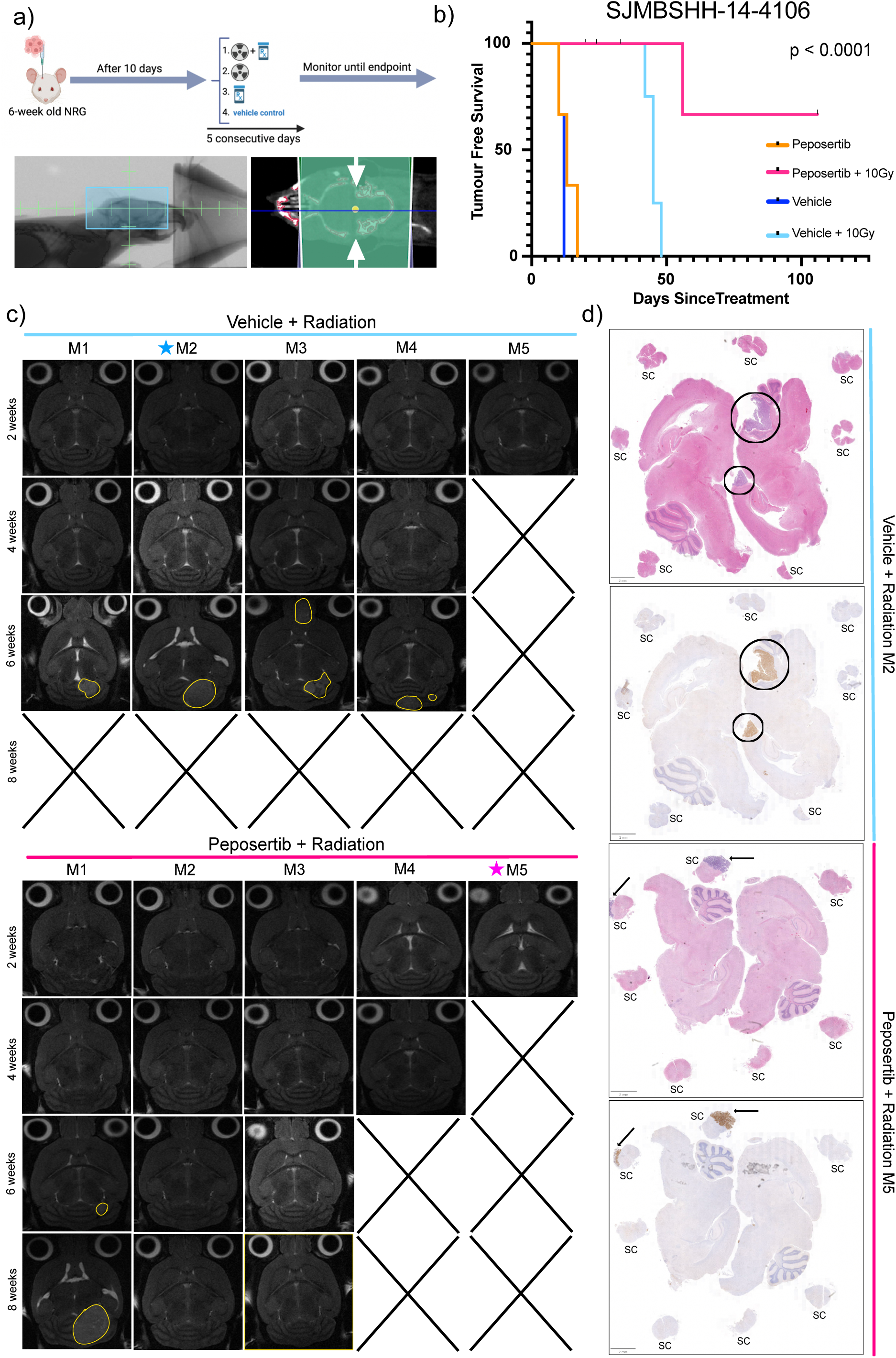
Inhibition of DNA-PK with Peposertib results in potent radiosensitization *in vivo*. **a)** Experimental schema (top). PDX models were orthotopically injected into the hindbrain of 6-week-old male NRG mice. Following 10 days post-intra-cranial injections, mice were divided into four treatment cohorts treated with peposertib with and without radiation for 5 consecutive days. Field of radiation (bottom) with 10×20mm rectangular collimator. 2Gy was delivered in a bilateral beam arrangement. **b)** Kaplan-Meier survival plot of *TP53* mutated SHH-MB PDX demonstrating a significant increase tumor-free survival (p<0.0001) when treated in combination with peposertib and radiation. **c)** MRI images of all mice treated with radiation alone (top) and with radiation and peposertib (bottom) following treatment. Yellow contouring highlights bulk tumor. **d)** H&E (purple) and ki67 IHC (brown) of brain and spinal cord (SC). Circles represent bulk tumors and arrows represent metastasis. Representative mice are highlighted in Figure 6c.

**Figure 7:**
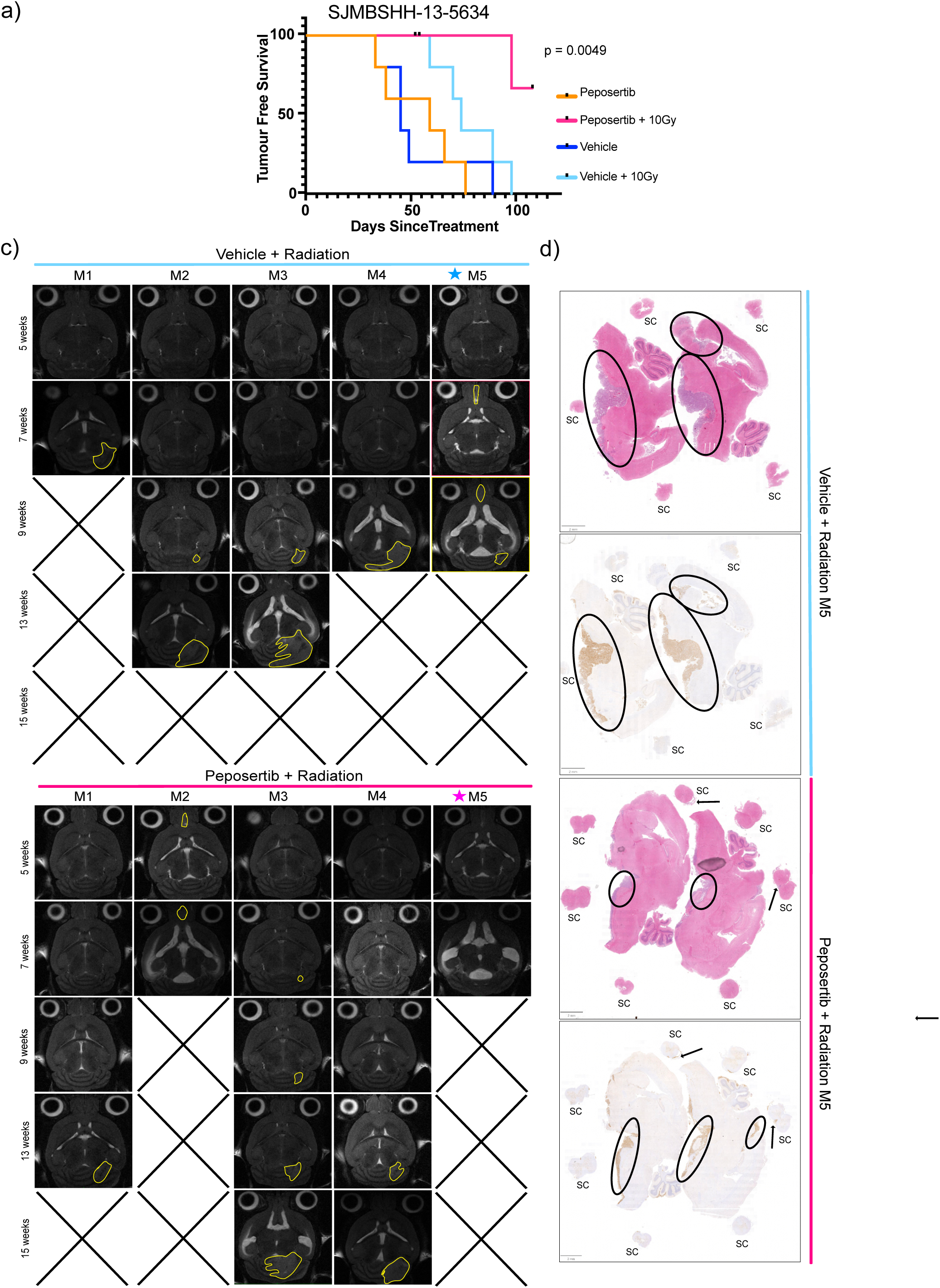
Inhibition of DNA-PK with Peposertib results in potent radiosensitization *in vivo*. **a)** Kaplan-Meier survival plot of *TP53* mutated SHH-MB PDX demonstrating a significant increase survival (p<0.0026) when treated in combination with peposertib and radiation. **b)** MRI images of all mice treated with radiation alone (top) and with radiation and peposertib (bottom) following treatment. Yellow contouring highlights bulk tumor. **c)** H&E (purple) and ki67 IHC (brown) of brain and spinal cord (SC). Circles represent bulk tumors and arrows represent metastasis. Representative mice are highlighted in Figure 7b.

## Discussion

Despite aggressive multimodal therapy, TP53-mutant SHH medulloblastoma remains one of the most challenging entities in pediatric cancer. Several strategies are employed worldwide, including current efforts to omit alkylator therapy, however, none have been successful so far in the clinic and survival remains dismal. Pre-clinically there has been a paucity of promising targets for *TP53*-mutant SHH-MB that significantly increase survival *in vivo*, and there have been a limited number of radiosensitizers identified with activity in bonafide SHH-MB systems.

Our finding using genome-wide positive selection screens that loss of *Trp53* as the main mediator of radioresistance in SHH-MB are consistent with previous studies using a sleeping-beauty transposon system combined with radiotherapy, and with human studies of relapsed medulloblastoma. Indeed in human studies sequencing primary and recurrent SHH-MB, *TP53* mutations emerge at relapse in a subset of relapsed tumors, and are conserved from diagnosis through recurrence, with a preponderance for local relapse in the high-dose radiation field. Overall the observation that *TP53* inactivation is both responsible for radiation resistance at diagnosis and at relapse support its role in being the major modulator of radiation failure in both murine and human SHH-MB. We also observe some sensitization of TP53 wild-type SHH-MB suggesting that the dose of radiotherapy may potentially be reduced for this group harboring a very favourable prognosis.

We devised an innovative genome-wide approach to identify synthetic lethal interactions with radiotherapy. Our finding that only components of non-homologous end-joining are synthetically lethal with radiotherapy is surprising as p53 has known roles in both homologous recombination and non-homologous end-joning, suggesting that this might represent a specific vulnerability in medulloblastoma. A limitation of our screen is the use of *in vitro* models of murine SHH-MB, due to the lack of bonafide human models of SHH-MB that propagate *in vitro*, however, our models display many features consistent with human *TP53*-mutant SHH-MB including chromosomal instability and significant radioresistance. Our observation that inhibition of DNA-PK, the prime mediator of non-homologous end-joining, as being a potent radiosensitizer in two bonafide PDX models of somatic and germline *TP53-*mutant SHH-MB, establish this as a promising treatment strategy that should be pursued in this very high-risk group.

Previous studies have eluded to using p53 status as a predictive biomarker for response to DNA-PK inhibition due to the provactive role p53 plays within the DDR. When treated with a combination of radiation and DNA-PK inhibition, tumors with normal p53 status increase their protective pathways through reinforced cell cycle arrest. In contrast, tumors with dysregulated p53 function accumulate genomic instability and result in cell death through mitotic catastrophe.^33,34^ Other studies have found consistent interactions between DNA-PK and p53 in determining radiation sensitivity.^35^

Peposertib is currently being evaluated in human clinical trials as a radiosensitizer in a variety of entities including head and neck cancers, and relapsed glioblastoma. A Phase I study of relapsed glioblastoma showed a very promising safety profile when combined with radiotherapy and further developments are likely (NCT04555577). Peposertib also appears to be relatively brain-penetrant based on our studies, and a pharmacokinetic study of brain metastasis suggests much higher accumulation of peposertib into tumor with a disrupted blood-brain barrier compared to tumor, where accumulation was substantial in the tumor core as compared to normal brain.^36,37^ We did not observe any obvious necrosis in our post-treatment necropsy of either focal or craniospinal irradiation treated animals, further supporting limited radiosensitization of normal brain. Our *in vivo* findings of a significant extension of survival with the combination of peposertib and radiotherapy support a model whereby peposertib has sufficient accumulation into medulloblastoma tumors, however further human studies are required to confirm this finding.

A concern is the potential for radiosensitization of normal tissues, however, highly focal radiation using proton radiotherapy can help mitigate this risk. Whether there is a role for peposertib in the treatment of other subgroups of medulloblastoma, including those where *TP53* emerges at relapse is an ongoing question, however, our observation that the introduction of a dominant-negative p53 allele confers resistance supports this notion. Currently, there is a paucity of novel pre-clinical therapeutic candidates for the treatment of p53-mutant SHH-MB, and our discovery that peposertib is a potent radiosensitizer in vivo of this high-risk group warrants further clinical investigation. Overall our findings support translation of peposertib in combination with radiotherapy into an early phase clinical trial for p53-mutant SHH-MB, including both germline (Li-Fraumeni syndrome) and somatic variants.

## Acknowledgements

A.D. is supported by a doctoral scholarship (CGS-D) from the Canadian Institute for Health Research, the Ontario Graduate Scholarship (OGS) and Flank Fletcher Memorial Fund. V.R. is supported by a Canada Research Chair in Pediatric Neuro-Oncology and operating funds from the Canadian Institutes for Health Research (PJT-488102 and PJT-400627), Rally Foundation for Childhood Cancer Research and Kids Join the Fight, Canadian Cancer Society Research Institute, the C.R. Younger Foundation and Brain Canada through the Canada Brain Research Fund with the financial support of Health Canada and Alvin Segal Family Foundation Future Leader in Canadian Brain Research. S. A. is supported by the Canadian Institutes of Health Research (PJT-169054).

## Author Contributions

Conceptualization A.D. and V.R.; Methodology A.D., G.M., S.A., V.R.; Formal Analysis A.D., G.M., V.R.; Investigation A.D., C.F.D., L.S., L.A., M.S., D.T., J.M., B.G.,S.S.; Resources R.M., R.A., A.A., F.C., C.N., P.A.N., P.D., U.T.,V.R.; Data Curation A.D., G.M., A.A., P.A.N., V.R.; Writing – Original Draft A.D. and V.R.; Writing – Review & Editing A.D., G.M., S.A., V.R.; Supervision P.A.N., U.T., P.D., S.A., V.R.; Project Administration V.R.; Funding Acquisition V.R.

The authors declare no competing interests

## Supplemental Information

Supplemental Figures: Figures S1-S7

Tables S1-S2.xlsx: Tables S1 and S2

Tables S3-S10.xlsx: Tables S3-S10

Tables S11-S16.xlsx: Tables S11-S16

## Supplemental Figure Legends

**Figure S1: p53 loss confers radioresistance in SHH medulloblastoma (Related to Figure 1)**

**a)** Precision-recall plots of the CRISPR-cas9 screens for three SHH*trp53* models and one SHHwt model across all treatment arms. Blue represents untreated, orange represents sublethal dose, and green represents a lethal dose of radiation.

**b)** An isogenic model of *Trp53* deficient SHH-MB was developed by introducing a dominant negative mutation to SHHwt, mimicking p53 deficiency and significantly increasing radiation resistance (p=0.016). Cell viability was measured using alamarBlue. Data was normalized to 0Gy and a positive control and analyzed by Mann-Whitney test.

**c)** Radiation dose-response curve with one *Trp53* null SHH-MB. Cell viability was measured using alamarBlue and normalized to 0Gy and a positive control (puromycin).

Radiation dose-response curve of human Daoy SHH-MB. Cell viability was measured as described above.

**d)** Copy number plots generated with 5X low pass WGS across four *Trp53* deficient SHH-MB and the isogenic pair.

**Figure S2: Lethal doses of radiation do not yield a consistent resistance mechanism in Trp53 deficient SHH-MB models (Related to Figure 2)**

**a-c)** NormZ scores across all screened SHH*trp53* models treated with 8Gy (left) and 12Gy (right) lethal dose of radiation. Genes highlighted in pink have a positive NormZ score below the FDR threshold of 0.1 (10%) indicating they confer resistance with the top three ranked genes listed. Full list of genes can be found in **tables S11-16**.

**Figure S3: Peposertib is an ideal brain-permeable radiosensitizer. (Related to Figure 3)**

**a)** 12-point dose-response curves with and without radiation of three different DNA-PK inhibitors in three SHH*trp53* models. Analysis of the AUC from a non-linear regression demonstrates CC-115 and BAY-8400 do not sensitize to radiation (p=0.23, p=0.22, respectively) across all three models.

**b)** 12-point dose-response curve of peposertib with and without radiation in SHHwt.

**c)** Caco-2 permeability assay across four compounds between the apical (AB) and basolateral (BA) chamber. Anything above the green line represents ideal permeability (left). VX-984 and AZD7648 have a high efflux ration (right) and are not ideal compounds. Anything below the green line have a low efflux ratio, including BAY-8400 and peposertib, and are ideal permeable compounds.

**d)** HLM (right) and MLM (left) metabolic stability assays demonstrates all compounds show ideal metabolic stability (above the red line) in both human and mice.

**Figure S4: Therapeutic inhibition of DNA-PK sustains radiation sensitization over several cell doublings. (related to Figure 4)**

**a)** Cell growth assays over 7 days with and without radiation in combination with 250nM of Peposertib. Cell images were taken by Incucyte from four different fields/well every 2 hours over 7 consecutive days to infer cell growth. Cell confluence plotted as percent of the confluence determined in the DMSO controls (set as 100%). Each graph represents combined means from 2-3 technical replicates per model.

**b)** Representative images (10× magnification) of **Figure 4a and b** of one SHH*trp53* model (left) and Daoy (right) from one field of view obtained from the Incucyte.

**Figure S5: Peposertib and radiation incudes an increase G2 cell-cyle arrest. (Related to Figure 5)**

**a)** Representative images of population gating for analysis of flow cytometry.

**b)** Representative images of PI cell cycle analysis in one SHH*trp53* model at multiple timepoints. Purple represents % G1 phase, yellow represents % S phase, and green represents % G2 phase. Cell cycle was analyzed using the Watson Pragmatic algorithm.

**c)** Combined analysis of one SHH*trp53* modelwith three independent technical and biological replicates of cell cycle analysis. Left graphs represent % cell population in S-phase at 48 hours (top) and 96 hours (bottom). Right graphs represent % cell population in G2-phase at 48 hours (top) and 96 hours (bottom). Data analyzed by Kruskal-Wallis test with Dunn’s test for multiple comparisons.

**Figure S6: Inhibition of DNA-PK with Peposertib is a safe and potent radiosensitizer *in vivo.* (Related to Figure 6 and 7)**

**a)** Weight loss across treatment all four treatment arms showing no significant differences between treatment groups. Weight loss was calculated by subtracting the weight on the final day of treatment from the starting weight on day one. Data analyzed by Kruskal-Wallis test with Dunn’s test for multiple comparisons.

**b)** MRI images (top row) following 1-2 weeks after treatment of all mice treated without radiation. White circles highlight bulk tumor. H&E (purple, middle row) and Ki67 IHC (brown, bottom row) of brain at endpoint of all mice treated without radiation. Yellow contouring highlights bulk tumor.

**c)** H&E (purple) and Ki67 IHC (brown) of brain and spinal cord (SC) of all mice treated with radiation. Circles represent bulk tumors and arrows represent metastasis.

**Figure S7 Inhibition of DNA-PK with Peposertib is a potent craniospinal radiosensitizer *in vivo.* (Related to Figures 6 and 7)**

**a)** Craniospinal Radation: Field of radiation with 10×20mm rectangular collimator (brain), C10 collimator (cervical spine), and 10×30mm rectangular collimator (thoracinc spine). 2Gy was delivered in a bilateral beam arrangement.

**b)** Kaplan-Meier survival plot of *TP53* mutated SHH-MB PDX treated with craniospinal radiation demonstrating a significant increase tumor-free survival (p=0.0016) when treated in combination with peposertib and radiation.

**c)** MRI images of all mice treated showing tumors 7 days before treatment with follow-up after treatment. Radiation cohorts were treated with craniospinal radiation. Yellow contouring highlights bulk tumor.

**d)** Weight loss across treatment all four treatment arms showing no significant differences between treatment groups. Weight loss was calculated by subtracting the weight on the final day of treatment from the starting weight on day one. Data analyzed by Kruskal-Wallis test with Dunn’s test for multiple comparisons.

**e)** H&E (purple) and Ki67 IHC (brown) of brain and spinal cord (SC) of mice treated with radiation. Circles represent bulk tumors.

**f)** MRI images (top row) following 5 weeks after treatment of mice treated without radiation. Yellow contouring highlights bulk tumor. H&E (purple, middle row) and Ki67 IHC (brown, bottom row) of brain at endpoint of all mice treated without radiation. Circles represent bulk tumors.

## Methods

### Cell Culture

C57BL/6 *Trp53+/-* and *Ptch1+/-* mice were crossed where sporadic tumors arise in the same anatomical location and exhibit histology seen in MB. Cell lines used for *in-vitro* assays were derived from transgenic murine models generated from tumors at endpoint in either germline *Ptch1+/-;Trp53+/+* or *Ptch1+/-;Trp53+/-* C57BL/6 mice. These tumors were dissected and dissociated by gentle pipetting in PBS followed by filtering through nylon filters, following by resuspension in serum-free neural stem cell media (SHH2, SHH3, SHH4). SHH*trp53* #156 and SHHwt #130 were generated by Ward et al. Cells were maintained in serum-free neural stem cell media containing 20 ng/mL epidermal growth factor (EGF) and 20 ng/mL basic fibroblast growth factor (FGF) in laminin-coated plates. For the generation of the isogenic model, *Ptch1+/-;Trp53+/+* cells were infected with an MSCV-p53^DD^-GFP retrovirus and maintained as described above.^38^ Daoy cells were maintiained in 10%FBS in DMEM.

### Low Pass Whole-Genome Sequencing

To identify genomic copy-number aberrations from mouse whole-genome sequencing (WGS) data, adapter trimming was performed using bbmap (v37.28) with k-mer based trimming (k=23, mink=11) and minimum read length filtering (minlen=30). Next, we aligned the trimmed reads to the mouse reference genome (mm10) using bwa (v0.17.7). Duplicate reads were removed, and base quality scores were recalibrated using GATK (v4.1.1), using known SNP and indel sites from the mouse genome. To call copy-number aberrations we used QDNAseq (R package v1.40.0) to process the BAM files, including read binning, bias correction, filtering, and normalization. Lastly, segmentation of the normalized copy numbers was performed, and results were exported in BED format for downstream visualization.

### Radiation Dose Response Curves

2000-4000 cells/well (cell-type dependent) were plated in lamanin-coated 96-well plates and left to adhere overnight. The following day, cells were treated with increasing doses of radiation usint Best Theratronics. 72-hours after treatment, cells were treated with 10% volume of AlamarBlue viability reagent and incubated for 2-3 hours. Fluorescence was measured with a plate reader and cell viability was inferred by fluorescent intensity. Data was normalized to 0Gy plates and a positive control.

### Whole Genome Crispr Cas-9 Screening

CRISPR-Cas9 screens were performed using the mouse Toronto knockout library (mTKO) containing 94 528 targeting gRNAs and 418 non-targeting gRNAs.^26^ Approximately 200 000 000 cells were bulk transduced at an MOI of 0.3 with the gRNA library, with the addition of 0.8ug/ml of polybrene, for a minimum of 200-fold coverage and seeded onto 40 lamanin-coated 10 cm dishes. 24 hours following transduction, media was replaced with media containing 1ug/ml of puromycin. Following 72 hours of puromycin selection, cells were combined and collected into two 50 000 000 T0 samples and stored in -80 for later processing. The remaining cells were seeded across 30 plates (2 million per plate) for 2-3 doublings to allow for sufficient genome editing. Cells were then divided into 3 replicates, each containing 20 million cells seeded across 10 plates. For radiation sensitivity and resistance screens, two additional treatment arms were divided into 3 replicates per experimental arm with 20 million cells seeded across 10 plates per replicate. Sensitivity and resistance arms were treated with an experimentally determined single sublethal dose and a lethal dose every 2-3 doubling for a total of 3 fractions, respectively. Cell numbers were maintained to support a minimum of 200-fold library coverage. At timepoints of approximately 7 (midpoint) and 14 (endpoint) doublings from T0, samples were collected in a minimum of 20 million cell aliquots (200-fold coverage) for gDNA extraction.^39^

### Genomic DNA Library Preparation and Sequencing

Genomic DNA was extracted from T0, midpoint, and endpoint frozen cell pellets using QiaAMP DNA Blood Maxi Kit following the manufacturer’s protocol, followed by ethanol precipitation and resuspension at a concentration of 400ng/uL. For each sample, 50ug of gDNA was amplified in twenty 50uL PCR reactions using the KAPA HiFi Master Mix and 1uM of each outer TKOv3 primer for 18 amplification cycles. PCR reactions were pooled and 5uL was used as a template for a subsequent PCR reaction to add TrueSeq adaptor sequences and unique combinations of i5 and i7 barcode sequences to each sample. These reactions were performed using KAPA HiFi Master Mix and 5uL of a 10uM stock of barcode primers for 14 amplification cycles. Barcoded PCR products were gel purified and submitted for next generation sequencing using Illumina NextSeq500. T0 was sequenced at a depth of 500-fold and all other samples were sequenced at a depth of 200-fold.

### CRISPR-Cas9 Gene Knockout Studies

For single gene knockout studies, individual gRNAs were ligated into a BsmBI digested pLentiGuide mCherry/GFP-2A-Puro. Approximately 3.5 million HEK293T cells were seeded on 10cm plates 24 hours prior to transfection with 6 ug of lentiviral vector, 3.6 ug of PsPAX2 plasmid and 2.4 ug of VSV-G plasmid with OptimMEM and X-tremeGENE 9. Viral media was harvested 48-72 hours post-transfection, centrifuged at 1000 x g for 5 minutes and filtered prior to concentrating with Lenti-X concentrator. Cells were then transduced with the above lentivirus and selected with 1ug/ml puromycin following 72 hours. Knockout was confirmed via western blots.

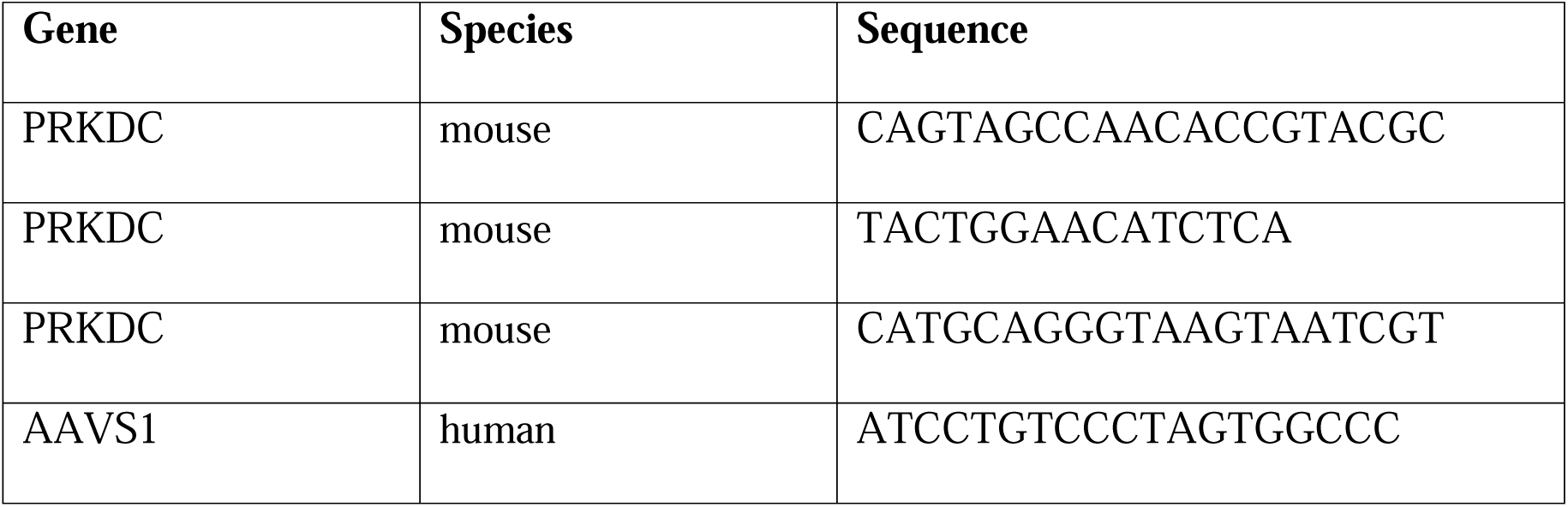

### Drug Dose Response Curves

Cells were plated at 2000-4000 cells/well (cell-type dependent) in lamanin-coated 96-well plates and left to adhere overnight. Cells were treated DNA damage inhibitors Peposertib (Selleck Chemicals, cat#S8586), AZD7648 (Selleck Chemicals, cat#S8843), CC-115 (Selleck Chemicals, cat#S7891), BAY-8400 (MedChemExpress, cat#HY-132293), AZD1390 (Selleck Chemicals, cat#S8680)) at a 12-point concentration curve ranging from 5000 nM to 5 nM using the Tecan D300e Digital Dispenser. 1-2 hours following treatment, cells were treated with 0Gy or a sublethal dose of radiation (2Gy or 4Gy). 72-hours after radiation, cells were treated with 10% volume of AlamarBlue viability reagent and incubated for 2-3 hours. Fluorescence was measured with a plate reader and cell viability was inferred by fluorescent intensity. Data was normalized per plate to a DMSO vehicle control and a positive control.

### Liver microsomal metabolic stability

In the Phase I analysis, test compounds were incubated at a final concentration of 1 µM, a level assumed to be well below Km values to ensure linear reaction conditions and prevent saturation. Working stocks were first diluted to 40.0 µM in 0.1 M potassium phosphate buffer (pH 7.4) before being added to the reaction vials. CD-1 mouse (male) or pooled human liver microsomes (Corning Gentest) were used at a final protein concentration of 0.5 mg/mL. Duplicate wells were prepared for each time point (0 and 60 minutes). The reactions were conducted at 37°C in an orbital shaker set to 175 rpm, with the DMSO concentration held constant at 0.1%. Each reaction had a final volume of 100 µL, including the addition of an NADPH-Regeneration Solution (NRS) mix (Corning Gentest), which contained glucose 6-phosphate dehydrogenase, NADP+, MgCl2, and glucose 6-phosphate. At the 60-minute time point, reactions were stopped by adding 200 µL (2 volumes) of ice-cold acetonitrile containing 0.5% formic acid and an internal standard. Samples were then centrifuged at 4,000 rpm for 10 minutes to remove debris and precipitated proteins. Around 150 µL of the supernatant was transferred to a new 96-well microplate for LC/MS analysis.

A narrow-window mass extraction LC-MS analysis was performed on all samples using a Waters Xevo quadrupole time-of-flight (QTof) mass spectrometer to determine the relative peak areas of the test compounds. The percent remaining was calculated using the formula:

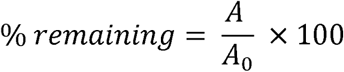

Where A is area response after incubation and A0 is area response at initial time point.

### Caco-2 Assay

Human epithelial Caco-2 cells (CRL-2102, C2BBe1) are seeded at a density of 40,000 cells per well on high-density PET membrane inserts with a 1.0 µm pore size and a surface area of 0.31 cm². These cells are used 21 or 22 days post-seeding, by which time the cell monolayers are fully polarized and differentiated. The permeability assay buffer consists of Hanks Balanced Salt Solution (HBSS) supplemented with 10 mM HEPES and 15 mM glucose, at a pH of 7.4. The dosing buffer includes 5 µM metoprolol (positive control), 5 µM atenolol (negative control), and 100 µM lucifer yellow, while the buffer in the receiver chamber contains 1% bovine serum albumin (BSA). The final concentration of the dosing solution is 5 µM in the assay buffer. Digoxin is used as a P-gp substrate control. Cell monolayers are dosed on either the apical side (A-to-B) or the basolateral side (B-to-A) and incubated at 37°C in a shaker set to 65 rpm. Samples for LC-MS analysis are collected from both the donor and receiver chambers at 120 minutes. Each determination is carried out in duplicate.

Narrow-window mass extraction LC-MS analysis is conducted for all samples using a Waters Xevo quadrupole time-of-flight (QTof) mass spectrometer to determine the relative peak areas of the parent compound. The percentage of transported drug is calculated based on these peak areas relative to the initial dosing concentration.

### Western Blotting

Western blotting was performed to investigate total and phosphorylated levels of H2AX and DNAPKcs protein utilizing specific antibodies. Cell lysate samples containing 20μg of protein were loaded next to a colorimetric protein ladder and separated by weight through sodium dodecyl sulfate-polyacrylamide gel electrophoresis (SDS-PAGE) on 12% polyacrylamide gels for ∼100 minutes at 110V. For DNA-PKcs, equal amounts of lysates were loaded onto NuPAGE 3-8% Tris-Acetate Gel. Samples were subsequently transferred to a 0.2μm PVDF membrane and blocking buffer (5% skim milk in TBST) was then added for 1 hour at room temperature. The PVDF membrane was incubated overnight with primary antibody at a temperature of 4°C, and the following day, after antibody removal and washing, the membrane was incubated with HRP-linked IgG anti-rabbit or anti-mouse secondary antibody for 1 hour at room temperature. Signal intensity was detected using the SuperSignal West Pico PLUS ECL substrate and visualized using a Licor Odyssey FC blot imager. Densitometry of the bands was performed with the Imagej Software. Data was normalized to protein loading control and untreated control sample.

### IncuCyte Live Cell Imaging Confluence and Cell-Death Assay

Cells were plated at 300 cells/well (0Gy plate) and 600 cells/well (2Gy or 4Gy plate) in lamanin-coated 96-well plates and left to adhere overnight. The following day, cells were treated with the experimentally determined IC50 of peposertib (250nM) for 1hr followed by 0Gy or a sublethal dose of radiation (2Gy or 4Gy). Real-time cell growth and cell death was monitored using the IncuCyte Live-Cell Imaging System (Essen Biosciences Inc). Both phase contrast and red fluorescent images were taken every 1.5 hours at 10 x magnification in 4 separate fields of view per well over 7 days. Real time growth was calculated as percent confluence and real time cell death was calculated by dividing the number of red objects by percent confluence.

### Clonogenic Assay

Cells were plated at 300 cells/well (0Gy) and 600-900 cells/well (2Gy or 4Gy) in lamanin-coated 6-well plates) in regular (0Gy) or conditioned (2Gy or 4Gy) media and left to adhere overnight. The following day cells were treated with 250 nM of peposertib or a vehicle control (DMSO) for 1hr followed by 0Gy or a sublethal dose of radiation (2Gy or 4Gy). Cells were left for 14 days following treatment with media top-up every 96 hours. At endpoint, colonies were fixed with glutaraldehyde (6.0% v/v) stained with crystal violet (0.5% w/v) for 30 minutes. Colonies were imaged by the Nikon smz25 dissection microscope and manually counted. Surviving fraction was calculated by dividing the number of colonies by number of cells seeded multiplied by plating efficiency using the equations below.

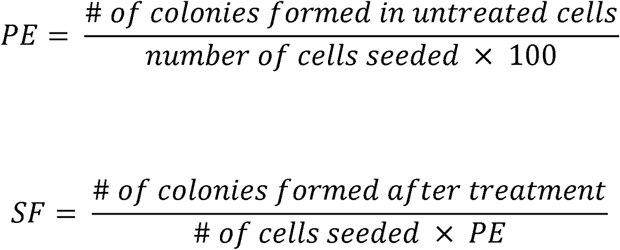

### Flow Cytometry

SHH*trp53* cells were treated DMSO, 250nM peposertib, 4Gy of radiation, or a combination of treatments over 48 and 96 hours. Cells were collected and resuspended in one million cells/ml per sample. Samples were fixed with either 4% PFA (senescence assay) or 70% cold ETOH (cell cycle analysis). Cells were spun down and washed with PBS. For the senescence assay, cells were incubated for 1hr at 37 °C without CO2 with 1:500 dilution of the probe for β-galactosidase hydrolysis followed by another PBS wash. Β-Gal signal was measured using a 488 nm excitation laser and 515/30nm bandpass filter. For cell cycle analysis, cells were spun down and washed with PBS followed by resuspension in 500uL of FxCycle Propidium Iodide (PI)/RNase staining solution. PI signal was measured using 561nm excitation laser and. 586/15nm bandpass filter. For Annexin/PI cell death assay, unfixed cells were resuspended in 100uL of 1× Annexin Binding Buffer and incubated for 15 minutes at room temperature with 5uL of Annexin V-FITC and 1uL of 100ug/ml propidium iodide working solution. 400uL of 1× Annexin Binding Buffer was added; samples were kept on ice until analysis. AnnexinV-FITC signal was measured using 488nm excitation laser and 515/30 bandpass filter; PI signal was measured using 561nm excitation laser and 610/20nm bandpass filter. Samples were acquired on a 5 laser FACSymphony A3 analyzer running FACSDiva9 software (BD Biosciences). Data analysis was performed using FlowJo software, version 10.10 (BD Biosciences).

### In vivo survival studies

NRG mice 5-6 weeks of age were orthotopically injected with 260 000 cells. Sex of the recipient mouse was matched to the sex of the PDX model. Following 10 days after injections mice divided into four treatment cohorts: vehicle alone, vehicle with radiation, peposertib alone, and peposertib with radiation. Vehicle was prepared as follows (300 mM sodium citrate buffer, 0.25% methylcellulose, 0.25% tween): 300 mM Sodium Citrate was prepared, pH 2.5. Methyl cellulose was added with constant mixing until the powder became completely dissolved. Lastly, Tween-20 was added. We allowed the solution to mix for 1-2 minutes to become completely homogenous and the final pH was adjusted to 2.5.

The peposertib treated cohort were treated with 50 mg/kg twice a day via oral a gavage. If treated in combination with radiation (5×2Gy), the first dose of peposertib was given 2 hours prior and 2 hours after radiation for five consecutive days. Image-guided radiation was delivered to the brain using the 10×20mm collimator and a bilateral beam arrangement. For craniospinal radiation, 2Gy was delivered in a bilateral beam arrangement to the brain, cervical spine, and thoracic spine using the 10×20mm, C10, and 10×30mm collimators, respectively. Mice were monitored with weekly or biweekly MRI’s. Humane endpoints were reached with 20% loss of maximum body weight or if clinically symptomatic. At endpoint, tumor cells were collected and whole brain and spine was formalin-fixed for IHC analysis.

### Immunohistochemistry

Brain and spine tissue were forlamin fixed (10%). Brain was cut saggitally and spine was cut axially placed is formalin-fixed, paraffin embedded (FFPE) blocks. FFPE blocks were thinly sliced, dewaxed, and stained with hetatoxykin and eosin or stained with the Ki67 antibody at a 1:600 dilution. Cancerous tissue was used as a positive control and normal heart and brain tissue is was used as a negative control when optimizing antibodies. Stained slides were imaged at 20× using the 3DHistech Slide Scanner. Images were visualized using QuPath0.5.1 software.

### Analysis of Genome-Wide Screen Data

Using MAGECK count function, read counts for individual gRNAs from triplicate samples were mapped to a gRNA library. The BAGEL algorithm was used to normalize gRNA reads to sequencing depth and to calculate the log2 fold-change relative to T0. Using a reference set of essential and non-essental genes, BAGEL computes a Bayes Factor (BF) of essentiality. To determine modifers to radiation resistance and sensitivity, DrugZ analysis was performed to calculate gene-level Z-scores. DrugZ normalizes read counts, calculates fold change and estimates the standard deviation for each fold change. It then provides a *Z*-score transformation and combines guide scores across replicates to a single gene score. Sensitizing genes were defined by a negative NormZ score within the 10% FDR threshold. Resistance genes were defined as a positive NormZ score within the 10% FDR threshold.

### Statistical Analysis

Statistical details of which tests were for each experiment can be found in figure legends. Unless otherwise stated, all statistical tests were completed with GraphPad Prism 9. All assays were conducted independitly, at least three time. All tests were run with at least n=3 biological replicates. P values ::: 0.05 were considered statistically significant. Unless otherwise stated, data represents mean and error bars represent SD. For in vivo orthotopic xenograft experiements, survival was measured using the Kaplan-Meier method with Log-Rank to compute statistical significance.

