## Supplementary figures and images for "Targeting synthetic lethality between non-homologous end joining and radiation in very high-risk SHH medulloblastoma"

### Supplemental Figures S1-S7

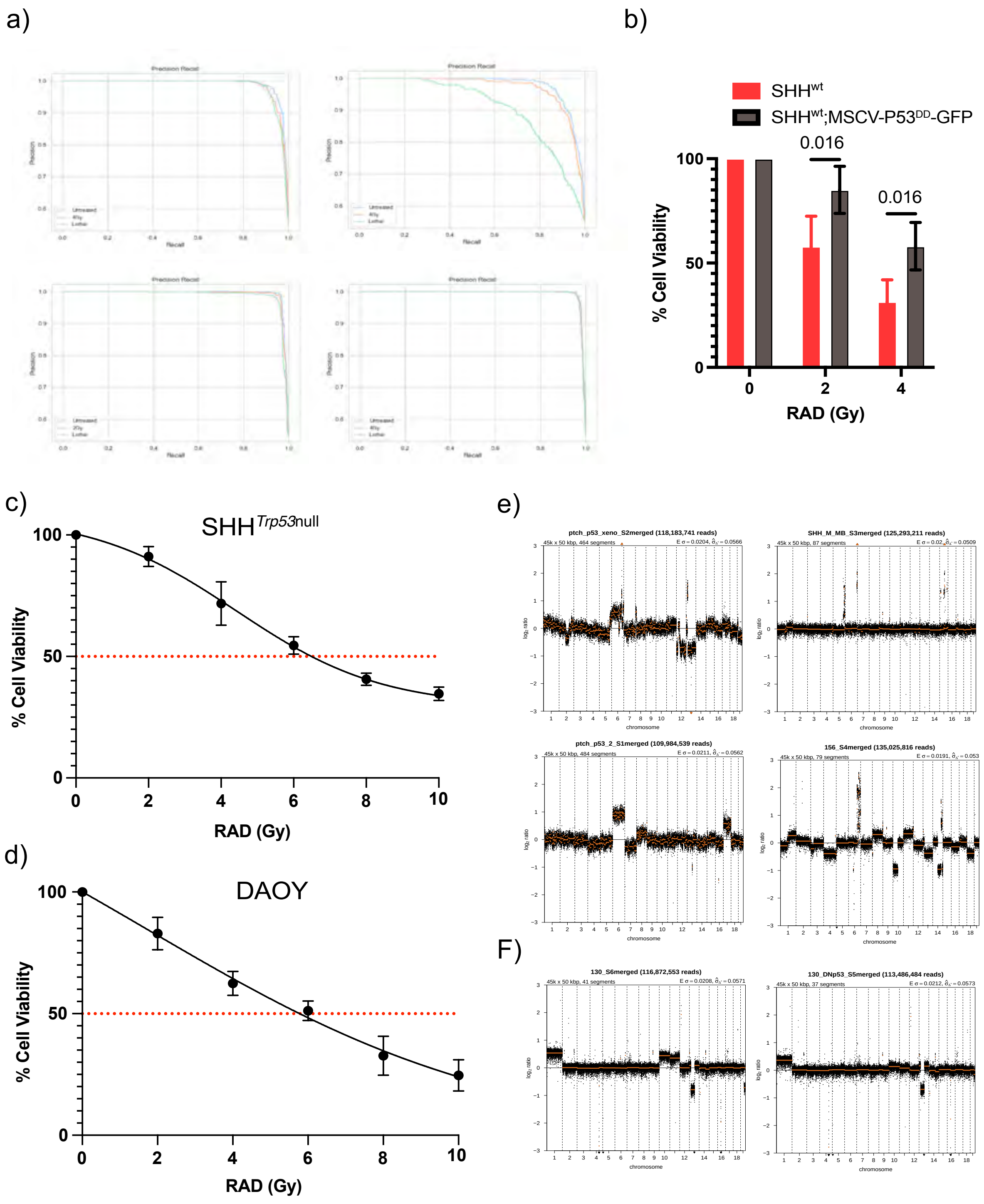

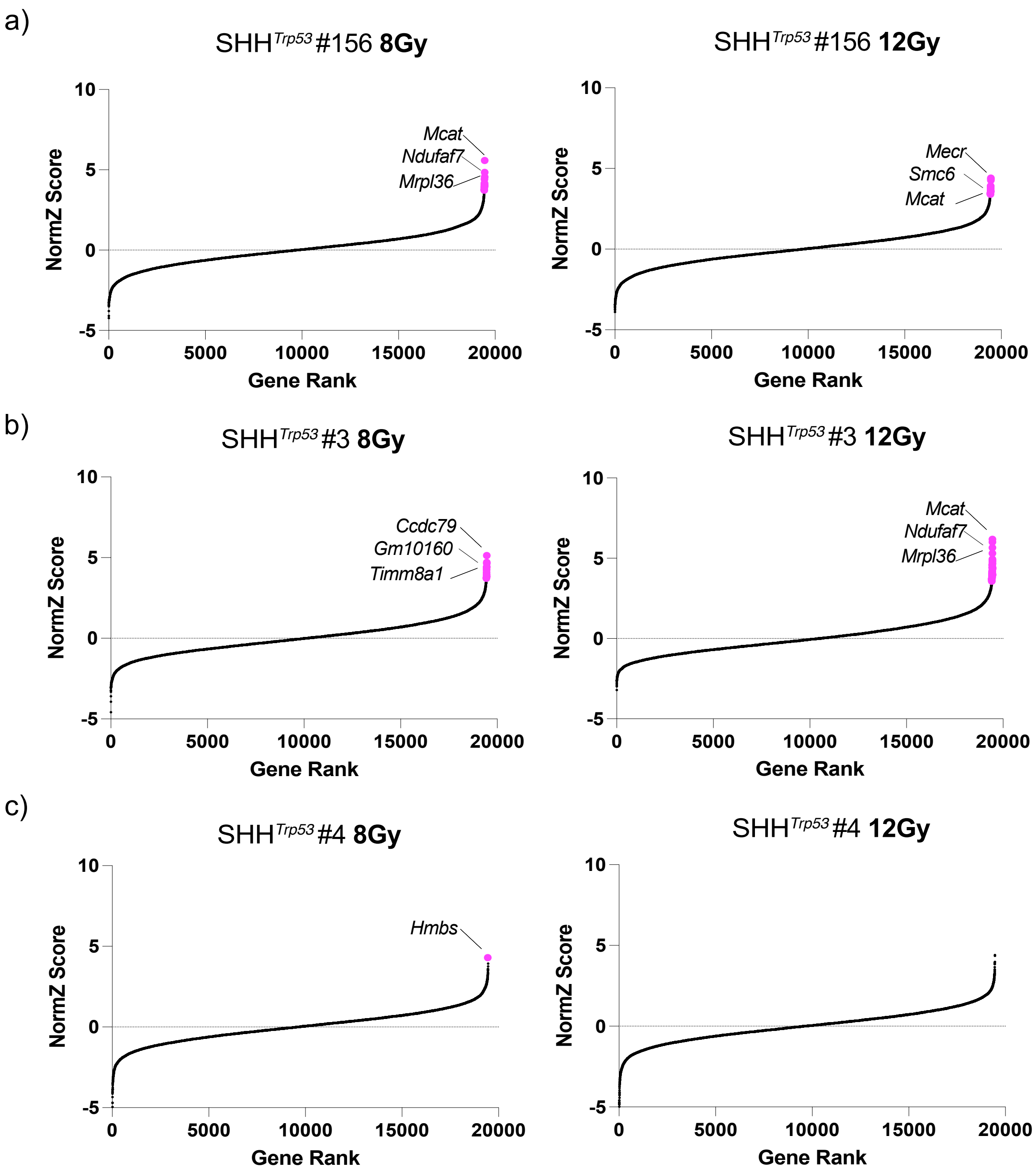

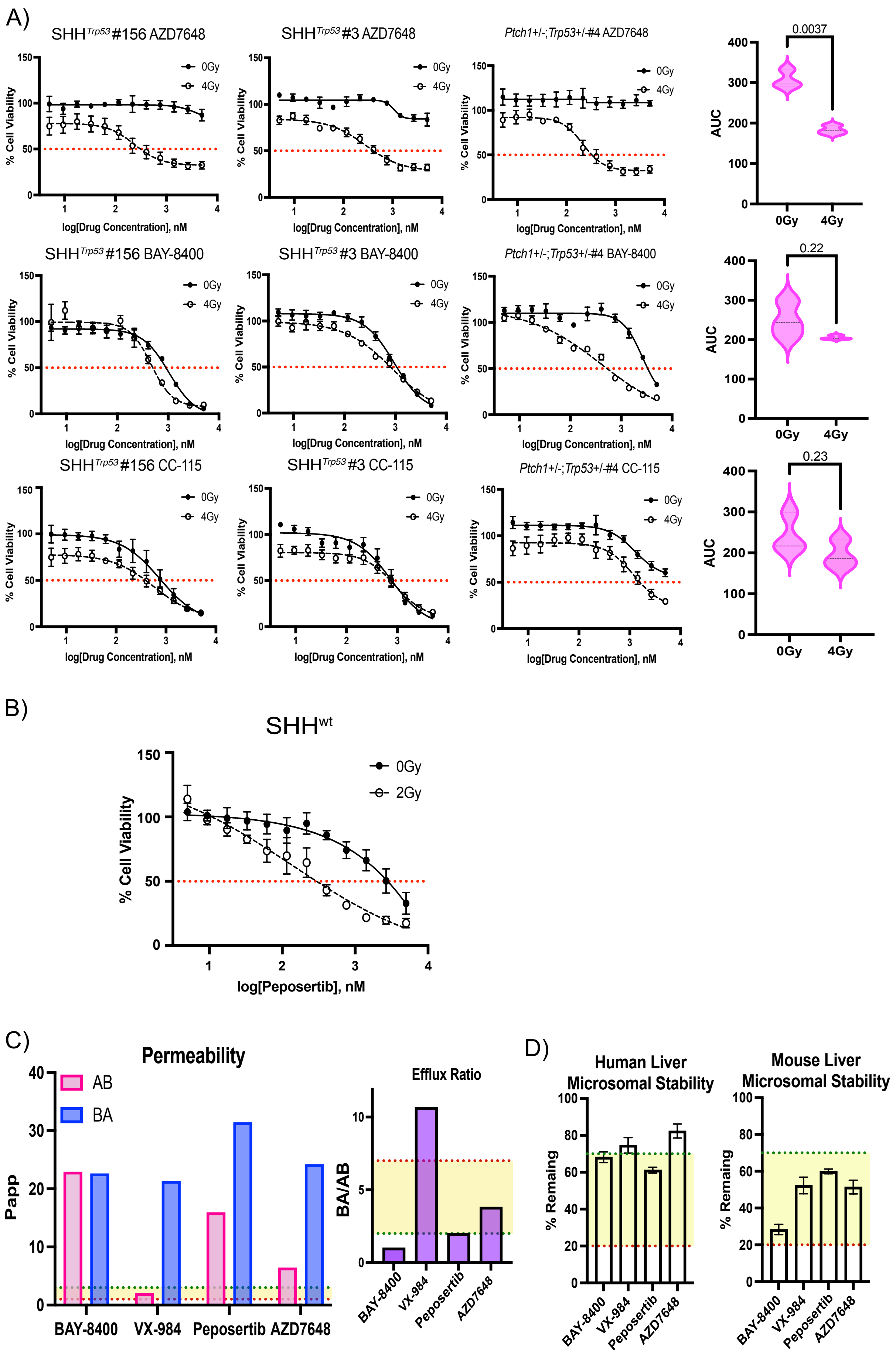

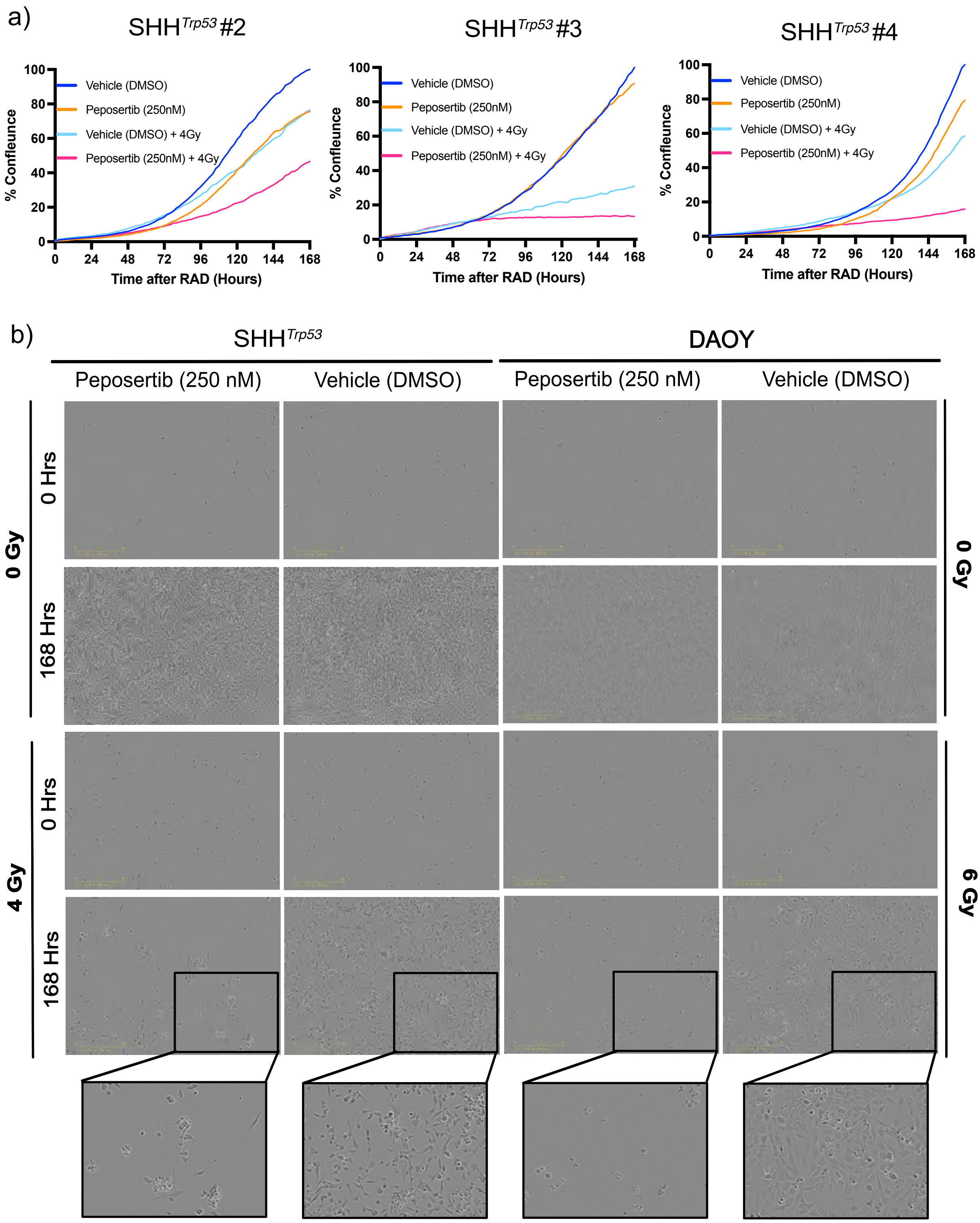

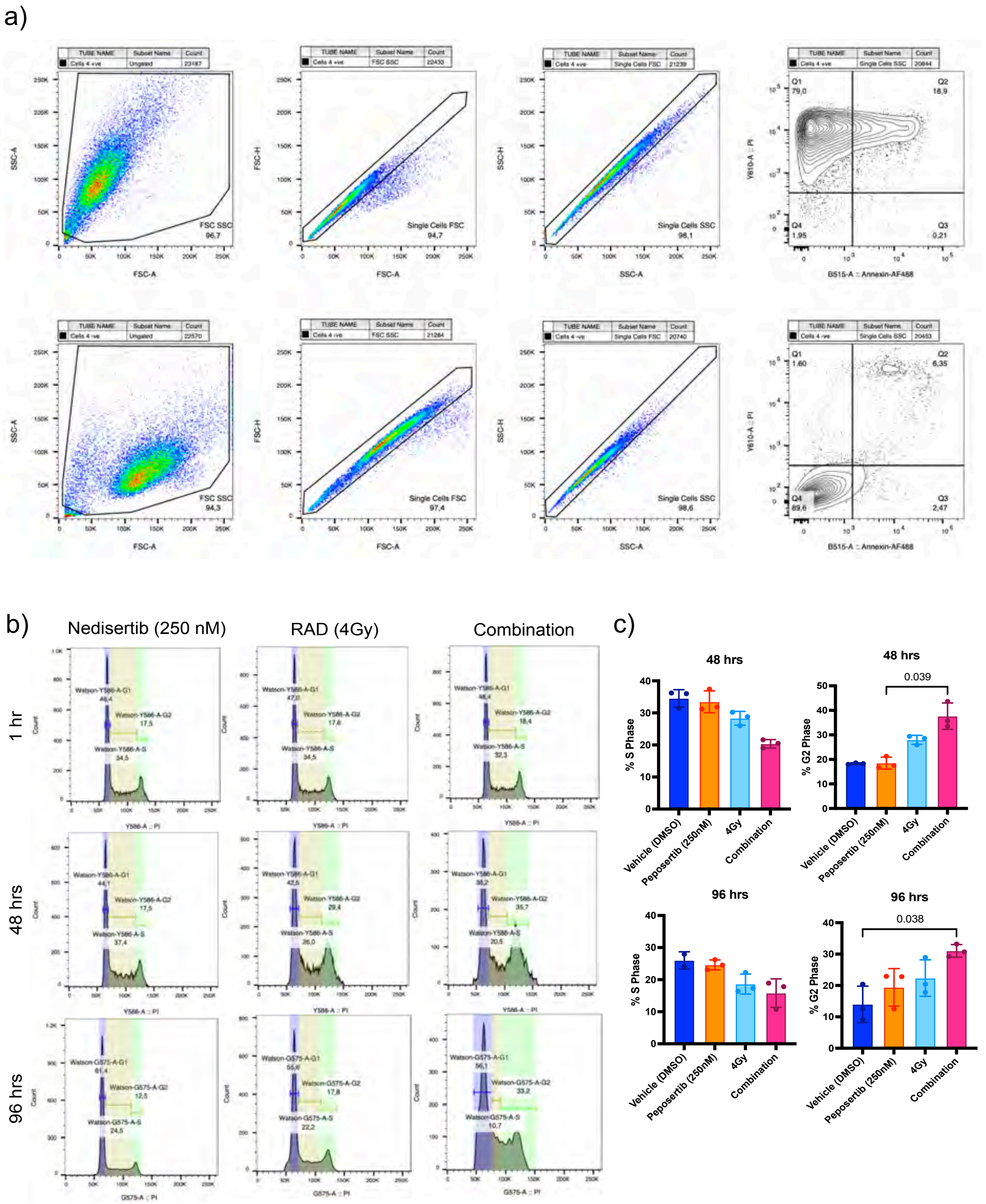

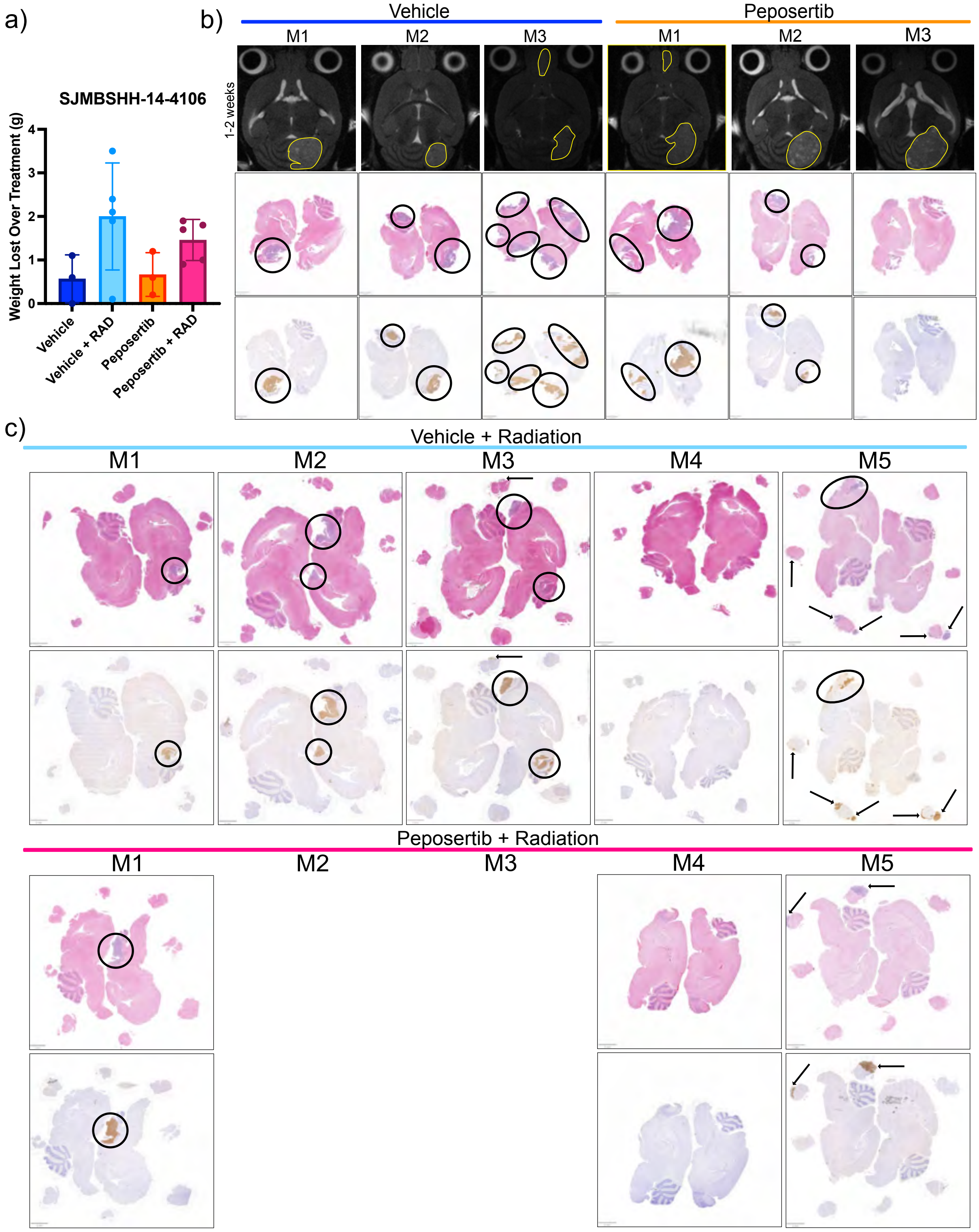

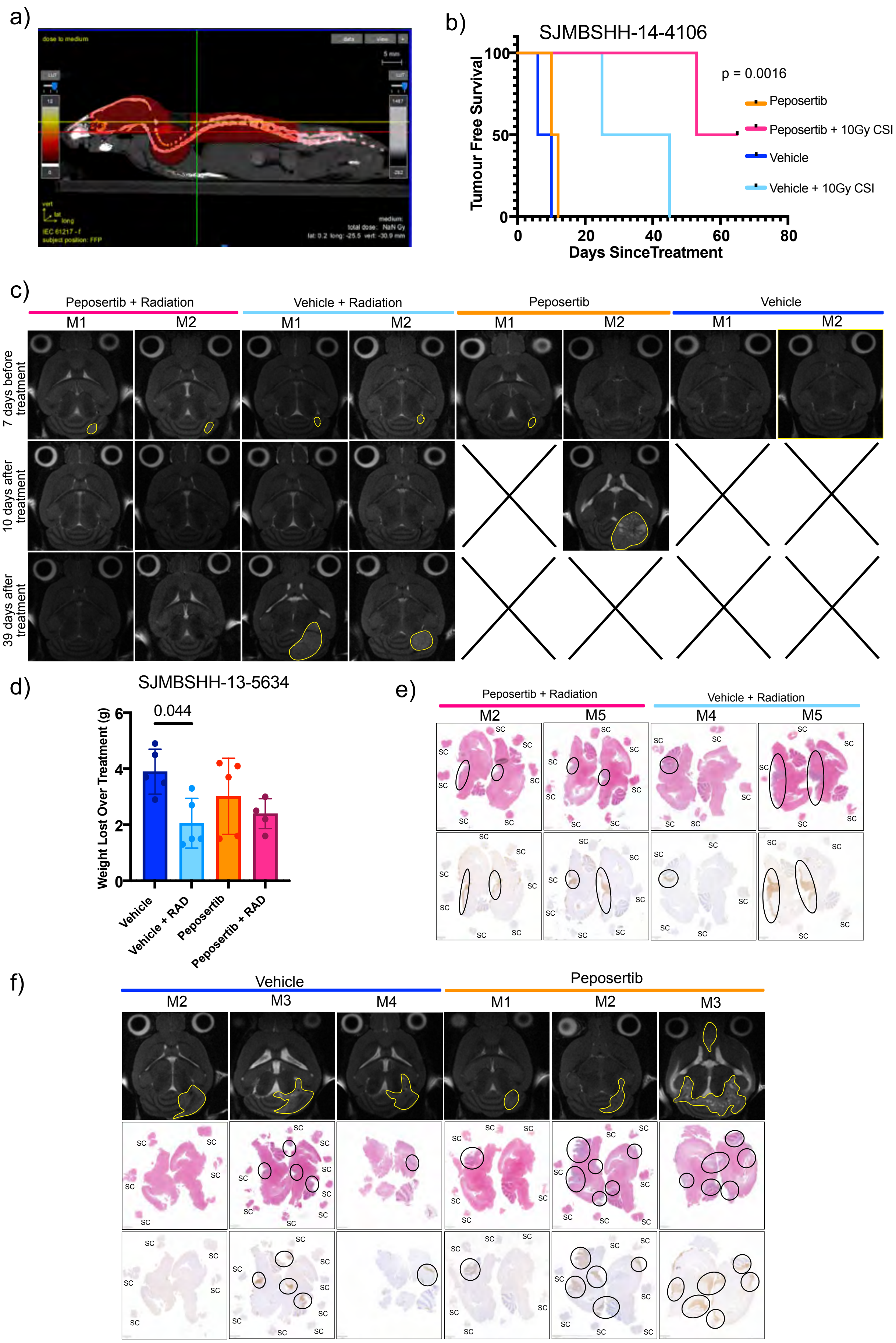
